# Single-cell RNA sequencing of peripheral blood mononuclear cells reveals complex cellular signalling signatures of metformin treatment type 2 diabetes mellitus

**DOI:** 10.1101/2024.01.04.574155

**Authors:** Jin-Dong Zhao, Zhao-Hui Fang

**Author notes:** Corresponding author: Zhao-Hui Fang, PhD, Chief Doctor, Department of Endocrinology, The First Affiliated Hospital of Anhui University of Chinese Medicine, No. 117 Meishan Road, Hefei 230031, Anhui Province, China.

## Abstract

**Objective:** Type 2 diabetes mellitus (T2DM) is a complex polygenic disease. The onset of the disease is related to autoimmunity. However, how immune cells function in the peripheral blood remains to be elucidated. Metformin is the first-line treatment. Exploring biomarkers of T2DM based on single-cell sequencing technology can provide new insights for the discovery of metformin treatment T2DM in molecular mechanisms.

**Methods:** We profiled 43,971 cells and 20,228 genes from peripheral blood mononuclear cells (PBMCs) of T2DM patients and healthy controls by single-nucleotide RNA sequencing.

**Results:** B cells, T cells, monocytes/macrophages, platelets, neutrophils, NK cells and cDC2s were grouped into 7 subclusters. Furthermore, T cells and monocytes/macrophages might be significantly correlated with the clinical characteristics of T2DM patients. RPL27 and AC018755.4 expression were strongly negative correlated with HbA1c. CD4+ T cells are mainly in the memory activation stage, and CD8+ T cells are effectors. The 50 genes whose expression varied with developmental time were associated with cytoplasmic translation, cell‒cell adhesion mediated by integrin, and the regulation of the inflammatory response. Monocytes/macrophages include classic monocytes and nonclassical monocytes. The GSEA results showed that the marker genes were enriched in the HALLMARK_INTERFERON_GAMMA_RESPONSE and HALLMARK_TNFA_SIGNALING_VIA_NFKB. The WGCNA results showed 14 modules. Meanwhile, TNFRSF1A is the most core genes in network interaction. Further analysis revealed ligand‒receptor pairs, including MIF-(CD74 + CD44), MIF-(CD74 + CXCR4), ANXA1-FPR1 and LGALS9-CD45.

**Conclusions:** Our study revealed that the transcriptional map of immune cells from PBMCs provided a framework for understanding the immune status of T2DM patients with metformin treatment via scRNA-seq analysis.

## INTRODUCTION

Diabetes mellitus is a metabolic disease characterized by hyperglycaemia whose cause is related to heredity, environmental factors, autoimmunity and other factors. Currently, diabetes is divided into four types, of which type 1 diabetes mellitus is mostly an autoimmune disease. In type 1 diabetes mellitus, T lymphocytes are activated in vivo, causing rapid destruction and the functional failure of islet beta cells, and the diseases eventually progresses to diabetes mellitus [1–2]. In China, type 2 diabetes mellitus (T2DM) is the most common form of diabetes, accounting for approximately 90-95% of cases, and its incidence is approximately 11.2%. T2DM comprises a group of heterogeneous diseases whose complex pathogenesis has not been fully elucidated [3]. The pathogenesis of T2DM mainly begins with insulin resistance, which leads to prediabetes and ultimately diabetes mellitus. Although there are many types of oral and injectable hypoglycaemic drug therapies, many patients still progress to having acute and chronic complications associated with diabetes, especially for vascular lesions.

Blood glucose is the main biomarker used for the diagnosis of T2DM. The discovery of other early biomarkers or molecular, pathological, and immunological changes is important for the diagnosis and evaluation of diabetes [4]. To date, the phenotypes and roles of T cells, NK cells, monocytes, macrophages and other immune cells have received less attention than other systems involved in diabetes mellitus [5–6]. In recent years, increasing evidence has shown that immune disorders are the main factors involved in the occurrence of T2DM [7–9]. Single-cell RNA sequencing (scRNA-seq) is a new technique that can be used to reveal cell heterogeneity and can be used to quantify the expression profile of each gene in each cell, which makes it easier to study genes. In addition to elucidating specific cellular functional alterations that may reveal cellular phenotypes and heterogeneity and providing a novel method for identifying biomarkers for the diagnosis and treatment of T2DM, these biomarkers may help predict the outcome of T2DM and associated complications [10]. The role of islet cell types in the genetic signalling pathways associated with T2DM susceptibility, particularly islet beta cell specificity, was identified by Rai V et al. [11–12]. Huang Y et al. identified six islet cell types and reported that SLC2A2, serinf1, RASGRP1 and CHL1 are biomarkers of T2DM; their analysis provided new markers for clinical diagnosis [13]. The 1 million cell atlas constructed by Wu H et al. reveals the types of kidney cells involved in the development of diabetic kidney disease. The identified DEGs can be used to explore underlying mechanisms as well as potential biomarkers for diagnosis [14]. Niu T et al. identified 11 cell types, and RLBP1 was found to be a therapeutic target for diabetic retinopathy [15]. Agrafioti P et al. reported that RELA was highly expressed and activated in the gingival macrophages of T2DM patients with periodontitis. This signalling cascade may be related to the pathogenesis and prognosis of periodontitis in T2DM patients [16].

In this study, we obtained scRNA-seq data from the plasma of healthy individuals and T2DM patients by labelling single-cell clusters and identifying key cell clusters using typical gene expression levels to understand the expression of every gene in every cell and the communication between cells. Metformin is the most basic treatment for T2DM due to its glucose-lowering effect. This analysis may provide new insights into the metformin treatment of T2DM.

## Results

### Comparison of clinical baseline information

There were no significant differences in age, sex, body mass index (BMI) or disease course between the two groups, but there were significant differences in fasting blood glucose and glucose levels between the two groups, as shown in Table 1.

**Table 1.** Comparison of clinical baseline information.

| Characteristic | Healthy group (n = 3) | T2DM group (n = 5) |
| --- | --- | --- |
| Age (y) | 56.33 ± 16.74 | 49.80 ± 2.95 |
| Sex (M/F) | 2/1 | 5/0 |
| Course of disease (y) | 0.00 ± 0.00 | 1.60 ± 1.29 |
| BMI (kg/m<sup>2</sup>) | 22.77 ± 2.57 | 25.46 ± 3.67 |
| FBG (mmol/L) | 5.37 ± 0.50 | 7.26 ± 0.37** |
| HbA1c (%) | 5.47 ± 0.25 | 6.56 ± 0.41* |

Note \*\* =P < 0.01, \* =P < 0.05, compared with the control group.
FBG: fasting blood glucose; BMI: body mass index; HbA1c: glycosylated haemoglobin A1c; FINS: fasting insulin; HOMA-IR: homeostatic model assessment of insulin resistance.

### Cell scRNA-seq

CD45+ cells were isolated from 8 blood samples for scRNA-seq. After filtration, there were 43971 total cells (2336-10146 per sample). Unsupervised clustering of single-cell data after normalization and aggregation using Seurat 3.0.2 identified seven cell types (Fig. 1A). The expression levels of genes in different types of cells are shown in a heatmap (Fig. 1B). We annotated these cells with classic gene markers, and the classic genes that were differentially expressed in these cells were consistent with the annotation (Fig. 1C). We found that the two groups of samples and the distributions of the seven cell types in each sample were different (Fig. 1D). The number of T cells and monocytes/macrophages in patients with T2DM significantly differed from that in the control group (Fig. 1E, P < 0.05).

**Figure 1:**
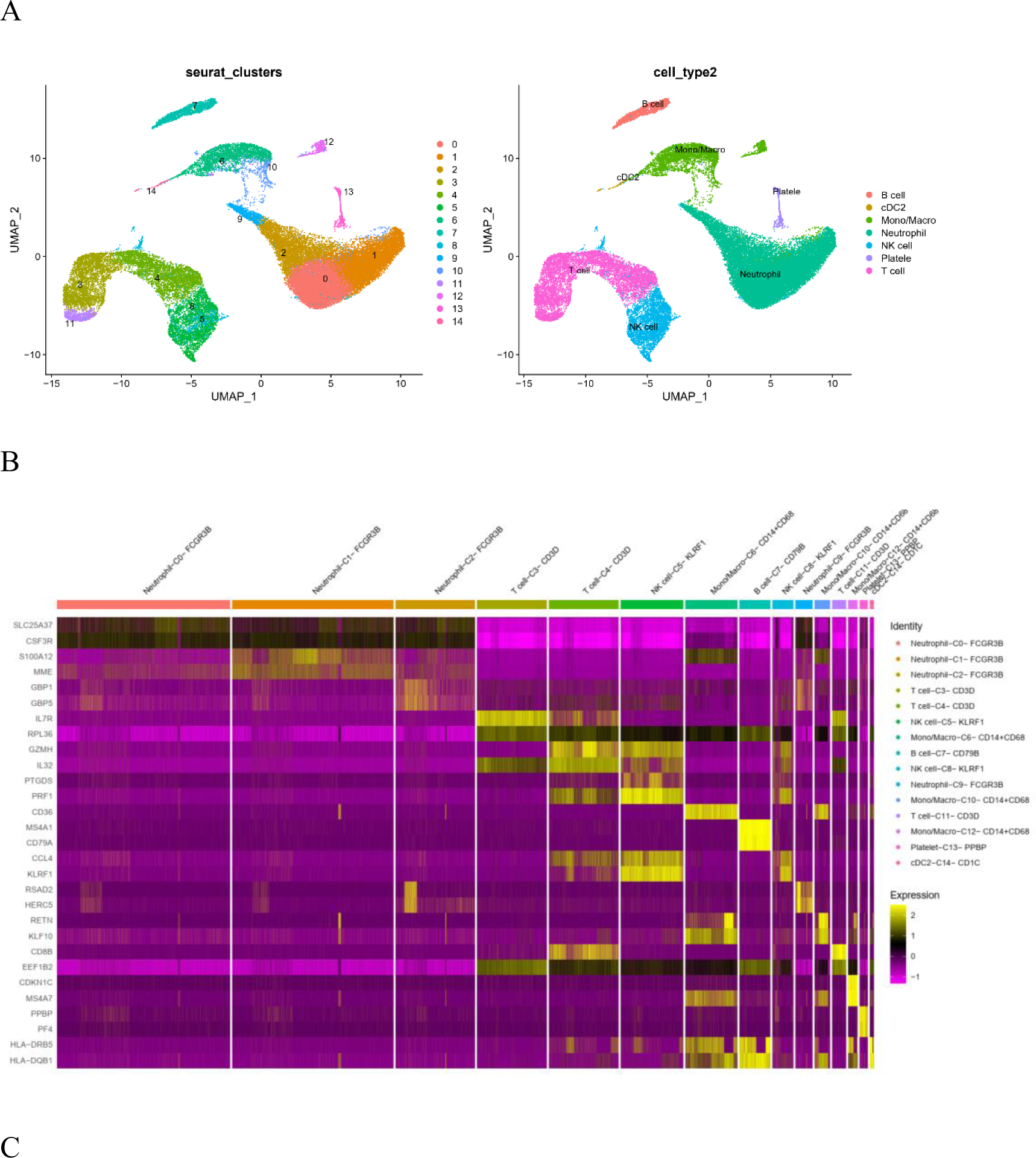
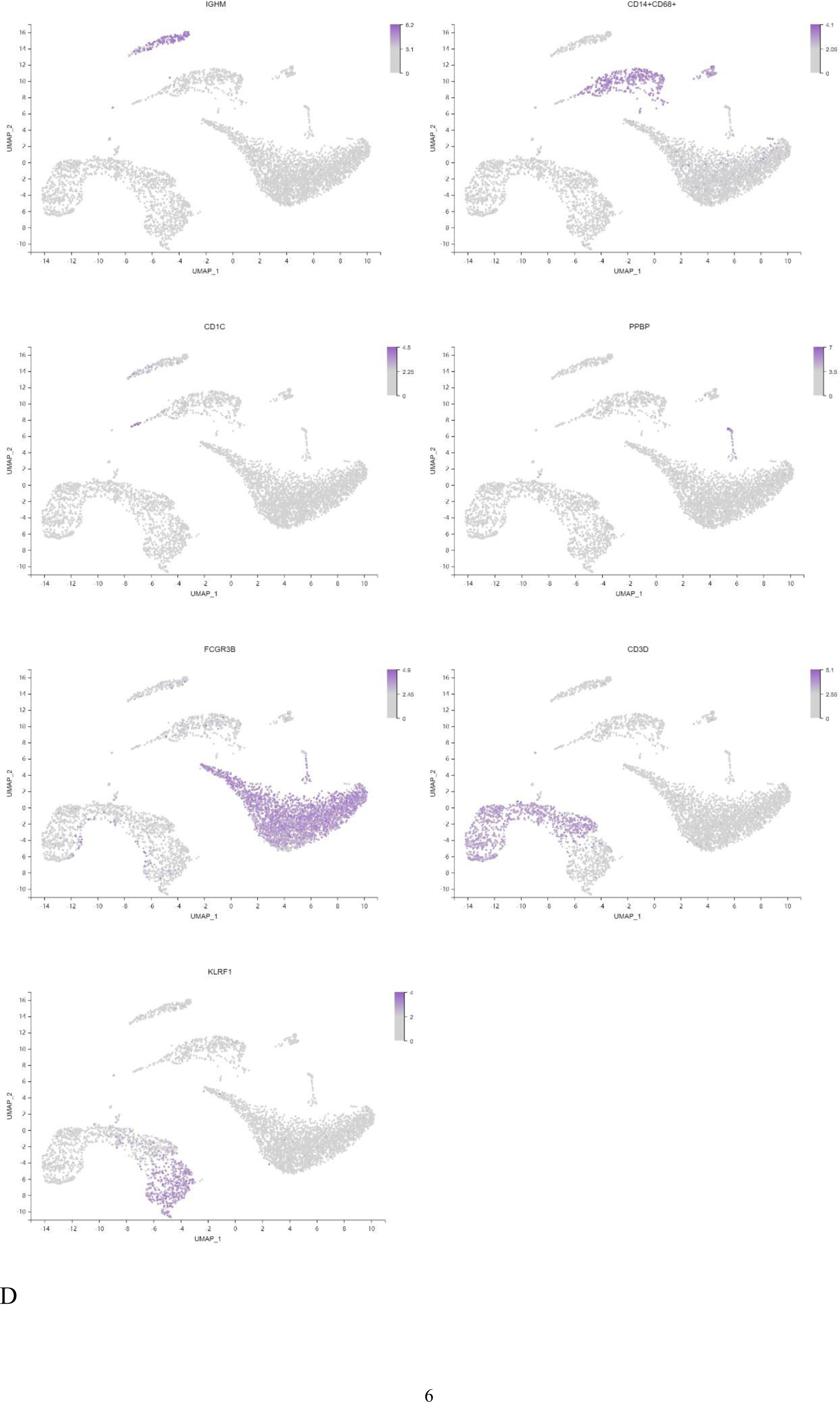
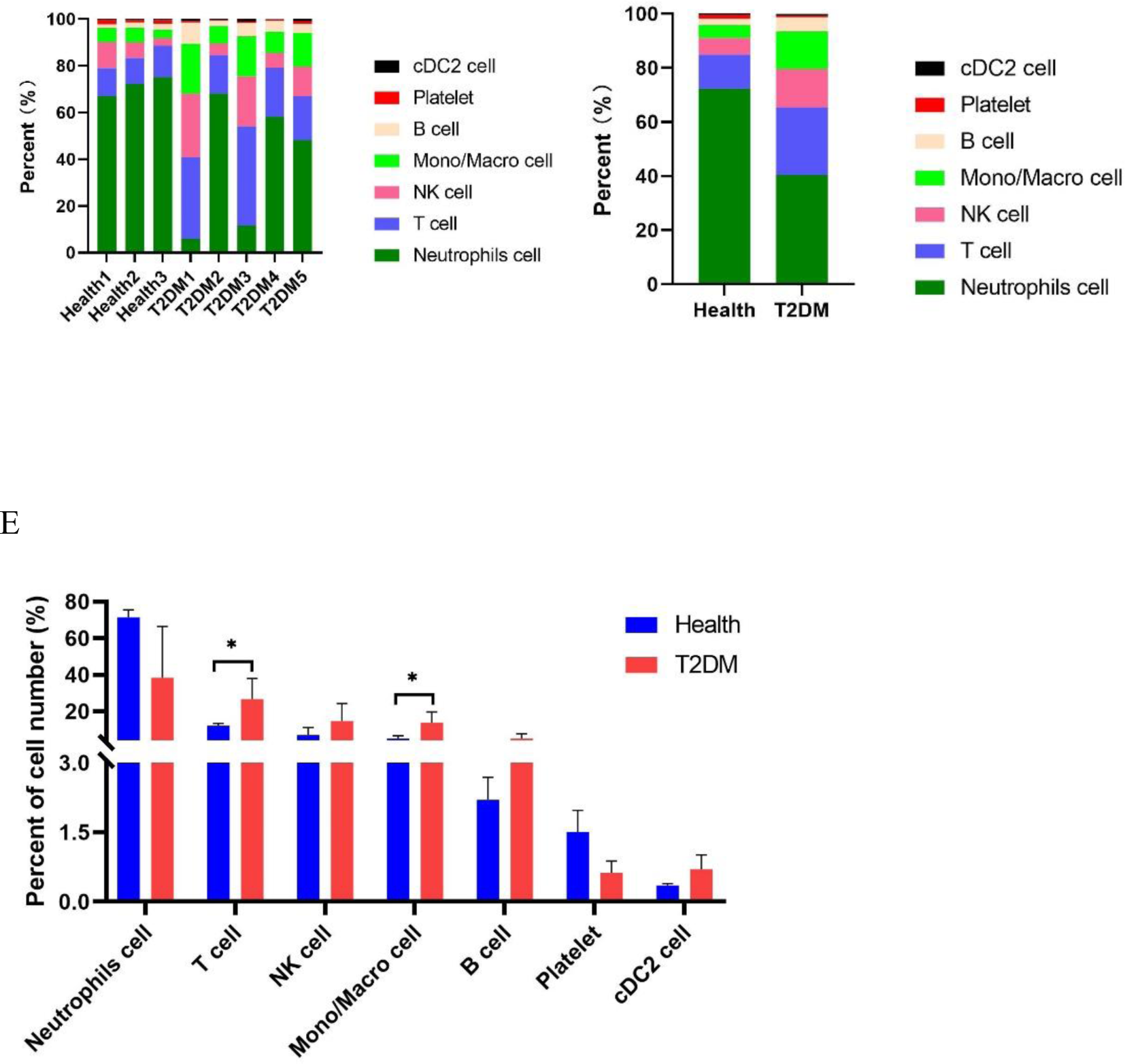
Cell atlas of immune infiltrates in PBMCs. A: UMAP plot of immune cell clusters. B: Heatmap of the relative expression levels of marker genes. C: Feature map showing the expression of the top marker genes for each cell type. E: Each sample corresponds to the cell types in each cluster. F: Two sets of samples were selected the identify the cell type component of each cluster. G: Comparison of the differences in the cell types of each cluster between the two groups of samples.

A total of 20228 genes were identified in the 8 samples. Among the 7 cell types, monocytes/macrophages showed expression of the most genes (Fig. 2A). Principal component analysis revealed that although the gene expression of the cells between the two groups was different, there was a certain degree of convergence (Fig. 2B). There were 3937 marker genes in the two groups that were enriched in KEGG pathways (Fig. 2C). Among these genes, 339 were related to endocrine and metabolic diseases. According to the KEGG pathway term level 2, 6 endocrine and metabolic diseases and 9 signal transduction pathways were screened (Fig. 2D). There were 11 marker genes associated with T2DM, which were enriched in KEGG pathways (Fig. 2E). The KEGG pathway relationship network corresponding to the 11 genes is shown in Fig. 2F. Notably, the GSEA results also identified glycolysis/gluconeogenesis (Fig. 2G). Correlation heatmap analysis was applied to evaluate the relationships between T2DM-related factors and representative genes. We found that MGST1 expression was most strongly positively correlated with FBG levels. The ANXA1, HK3, HK1, PKM and INSR genes were weakly correlated (Fig. 2H).

**Figure 2:**
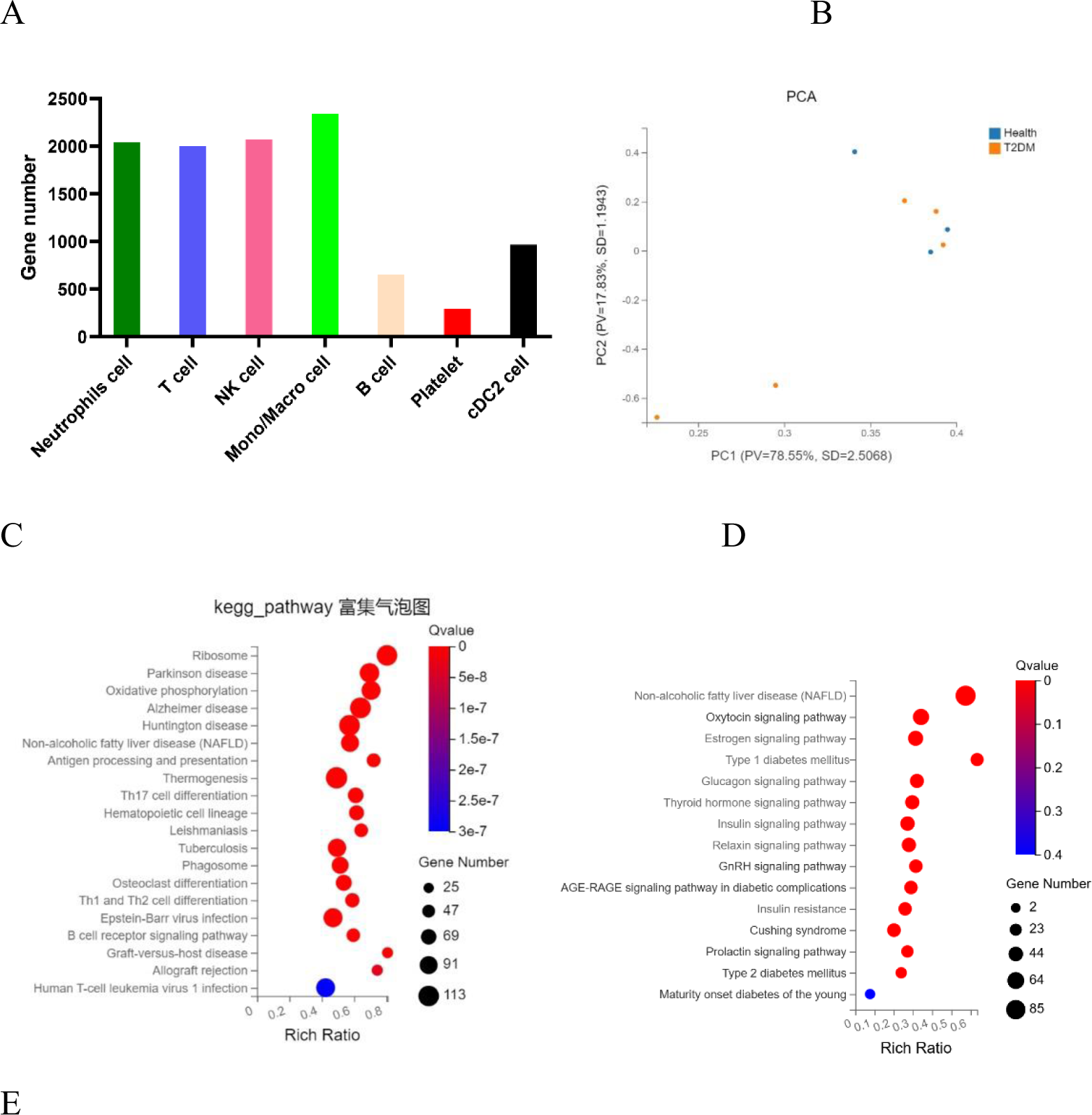
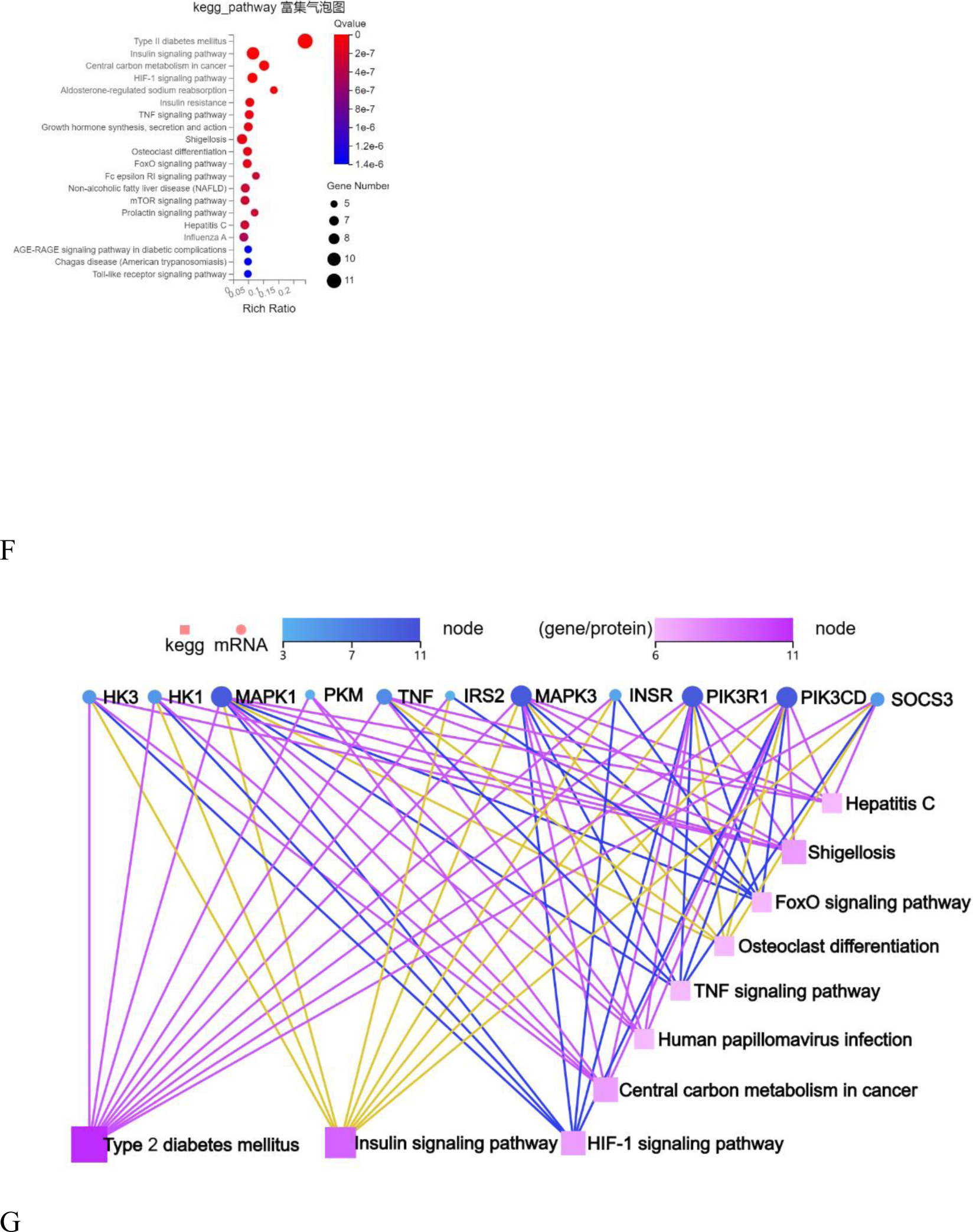
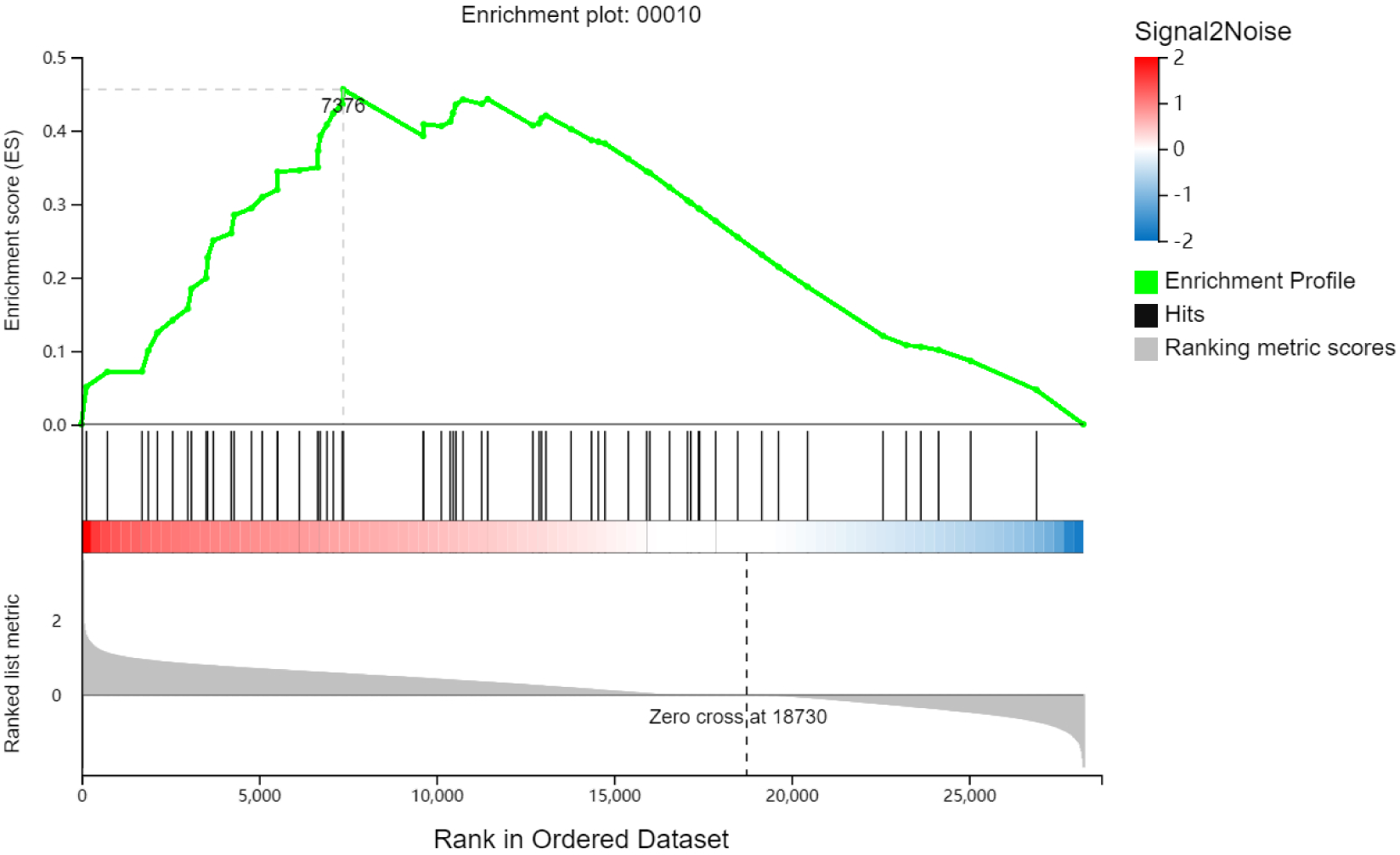
Map of PBMC genes and marker genes in T2DM patients. A: The number of genes in each cluster. B: PCA of genes in the two groups of cells. C: The KEGG pathways that were enriched by marker genes. D: KEGG pathways enriched in marker genes of the endocrine system. E: KEGG pathways enriched in marker genes of T2DM. F: KEGG pathways enriched in marker genes of the network corresponding to T2DM. G: GSEA of PBMCs.

### Clustering and subtype analysis of T cells and NK cells

T cells and NK cells were the main specific immune cells found in patients with T2DM, with a total cell count of 10965. Unsupervised clustering of T cells and NK cells identified two CD4+ T-cell clusters (including 3549 cells), two CD8+ T-cell clusters (including 3736 cells) and two NK-cell clusters (including 3680 cells) (Fig 3A). The expression levels of genes in different cell types are shown in a heatmap (Fig. 3B). T cells and NK cells were annotated separately by canonical genes, and the expression of the canonical genes of these cell types was consistent with the annotation (Fig 3B). There were 263 marker genes found in CD4+ T cells, 133 in CD8+ T cells and 626 in NK cells. The heatmap shows the average expression levels and percentage expression of representative marker genes in cells for healthy subjects and T2DM patients (Fig. 3C).

**Figure 3.**
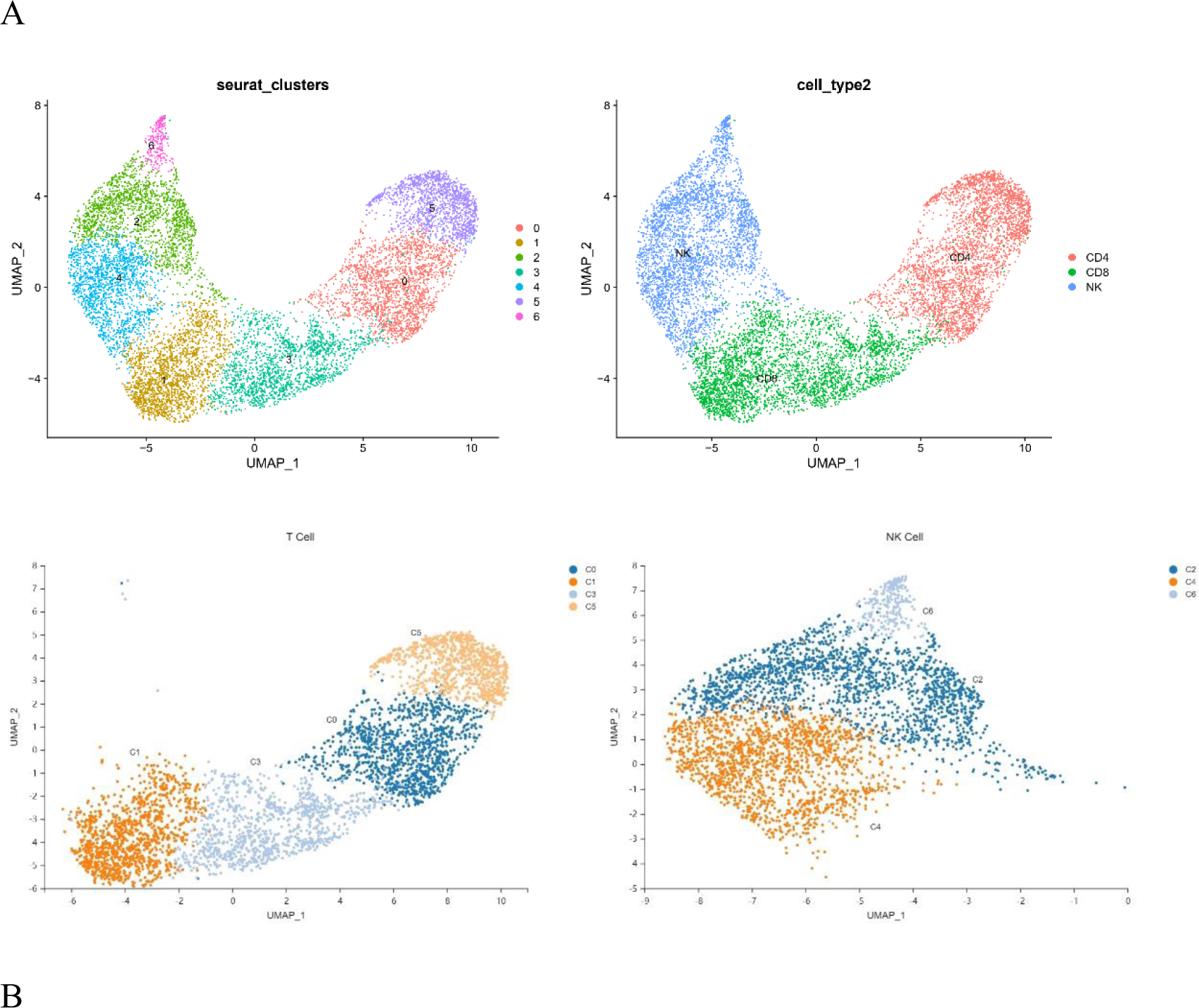
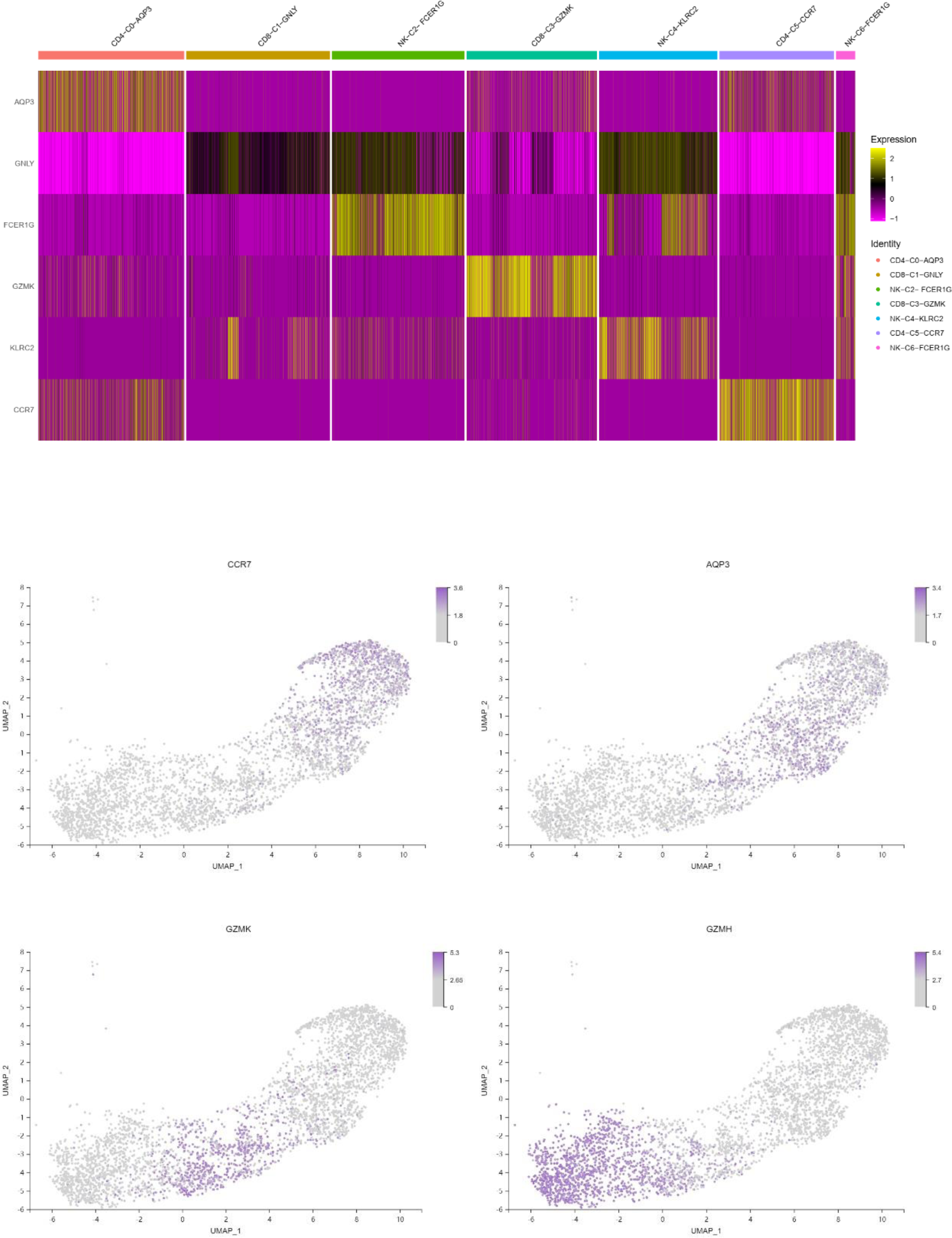
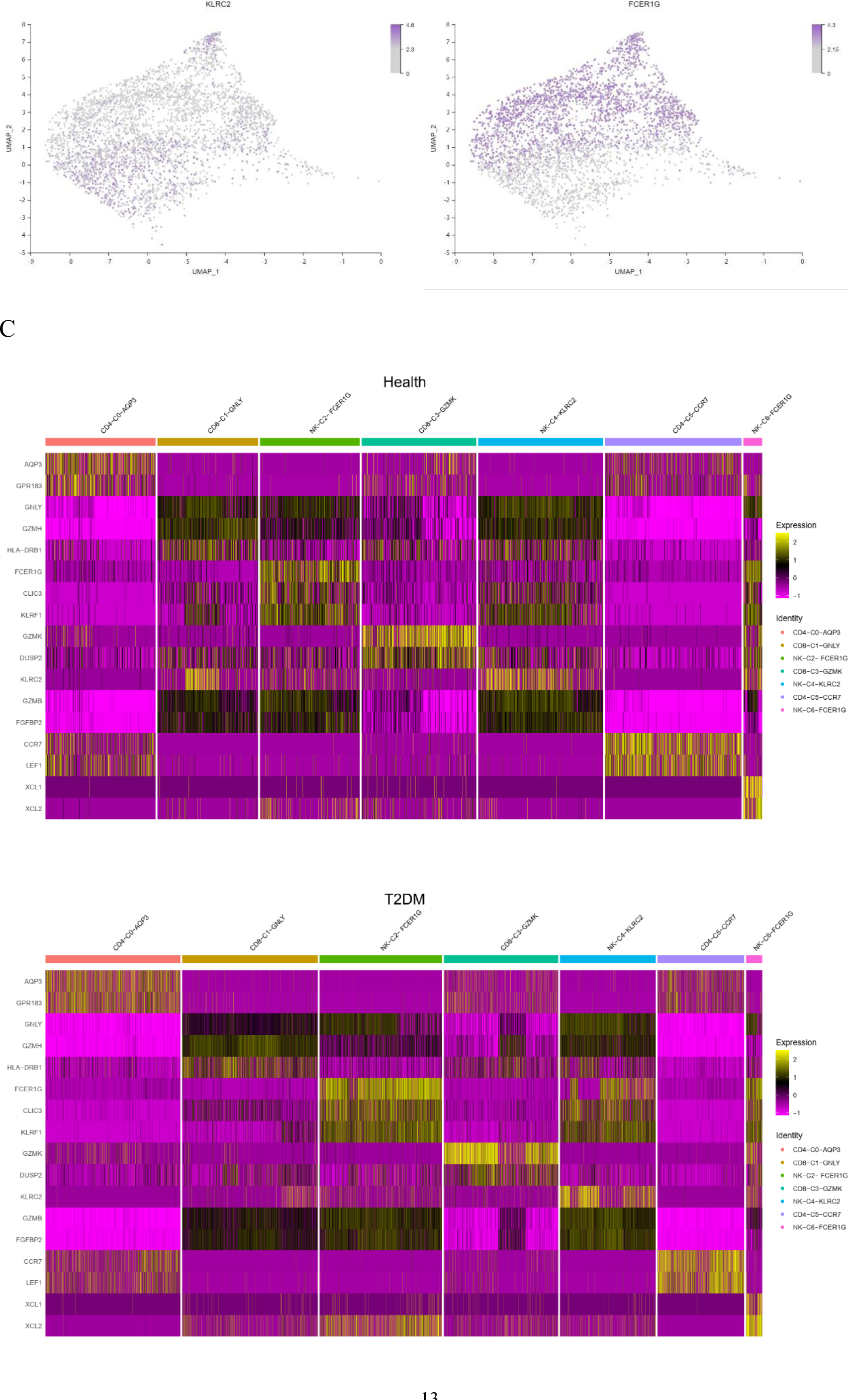
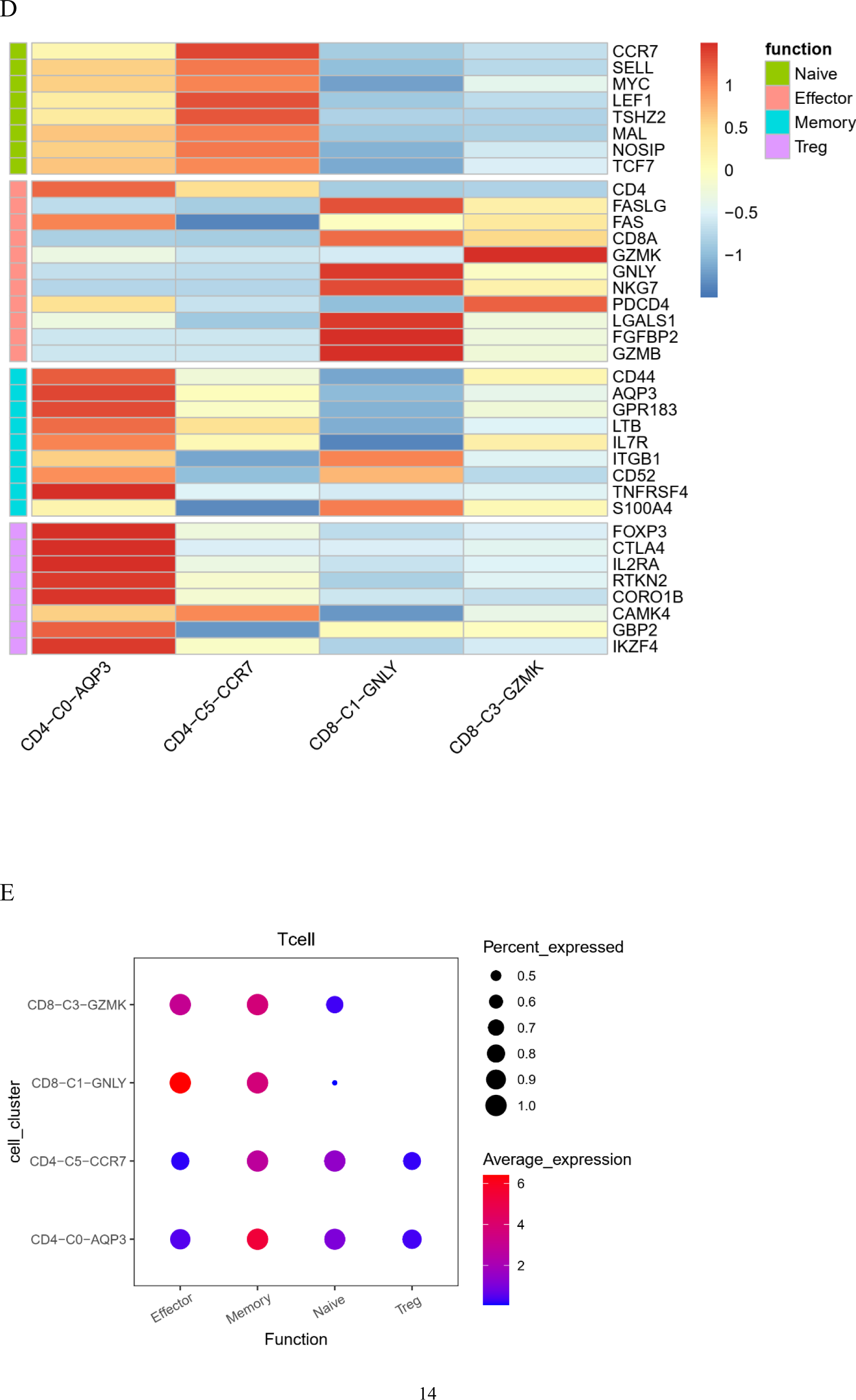
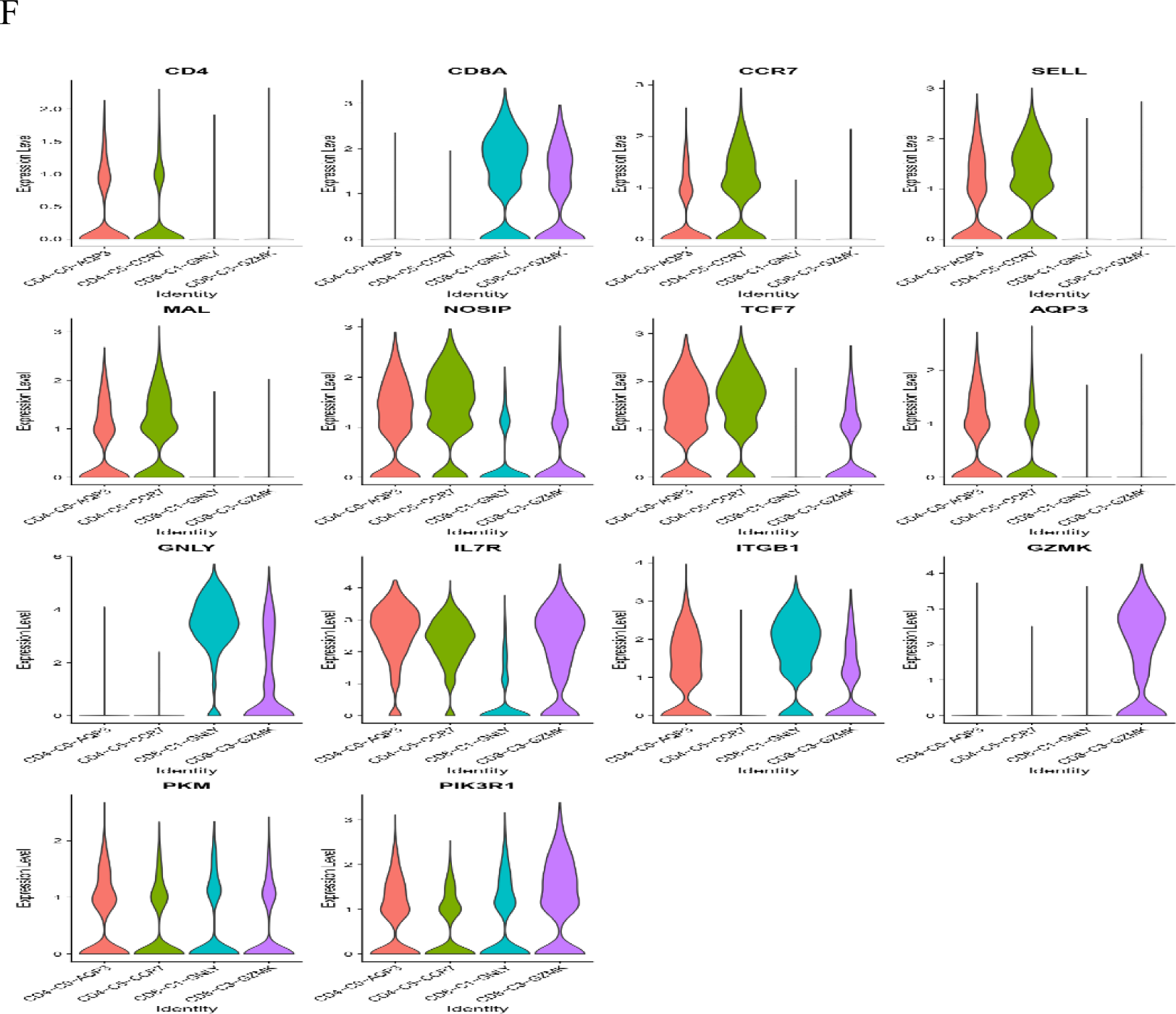
Characterization of T and NK cells. A: Uniform Manifold Approximation and Projection (UMAP) plot of T and NK cells. Four cell clusters were found among the T cells. Two cell clusters were found among the NK cells. B: A feature map showing the expression of marker genes for various cell types. The cells were identified by marker genes. C: Dot plot of representative signatures in T-cell clusters for healthy controls and T2DM patients. D: Heatmap of the expression of selected T-cell function-associated genes. E: Dot plot of representative activation stage signatures in T-cell clusters. F: Violin plot showing representative genes in T cells. The differences in the expression levels of genes in the T2DM group and healthy control group were compared to construct a volcano plot, as shown in Fig. 4A. Among these genes, 58 had upregulated expression and 61 had downregulated expression in T cells. We have conducted the correlation analysis between differentially expressed genes and clinical character. The RPL27, TXN1P and RPL37 of genes were negatively correlated with HbA1c. The MNDA of genes was negatively correlated with FBG, DDX5 was positive correlated. GIMAP7 was positive correlated with HOMA-IR (Fig. 4B). The GO enrichment results showed that the biological processes (BP) terms were mainly related to leukocyte chemotaxis, cytoplasmic translation, positive regulation of the apoptotic signalling pathway, myeloid cell activation involved in the immune response, the T-cell receptor signalling pathway, and the immune response-regulating signalling pathway. The cellular components (CCs) were mainly associated with cytosolic large ribosomal subunits, ribosomes, cytosolic ribosomes, tertiary granule membranes, and cell-substrate junctions. The molecular functions (MFs) were specifically related to structural constituent of ribosomes, Toll−like receptor binding, RAGE receptor binding, phospholipase inhibitor activity, and cell‒cell adhesion mediator activity (Fig. 4C). The KEGG pathway analysis results showed that the marker genes were enriched in the IL-17 signalling pathway, ribosomes, cell adhesion molecules, leukocyte transendothelial migration, and primary immunodeficiency (Fig. 4D). The first gene set, HALLMARK_TNFA_SIGNALING_VIA_NFKB, was selected to be used in generating a GSEA graph via HALLMARK pathway enrichment analysis (Fig. 4D). We further analysed the cell cycle progression of T cells in healthy controls and T2DM patients. In healthy controls, T cells remained in the G1, S and G2M states, and T cells in T2DM patients were in a proliferative state compared with those in healthy controls (Fig. 4E). The results of WGCNA showed that genes were divided into 7 modules. According to the correlation analysis between clinical character and the 7 modules, FINS and HOMA-IR were significantly positive correlated with module turquoise (Fig. 4F). The KEGG result of genes in the module turquoise is involved in the TNF signaling pathway, T cell receptor signaling pathway and NF-kappa B signaling pathway (Fig. 4G). Meanwhile, TNFRSF1A is the most core genes. We further analysed the cell cycle stages of T cells in healthy controls and T2DM patients. In healthy controls, T cells remained in the G1, S and G2M states, and T cells in T2DM patients were in a proliferative state compared with those in healthy controls (Fig. 4H).

**Figure 4:**
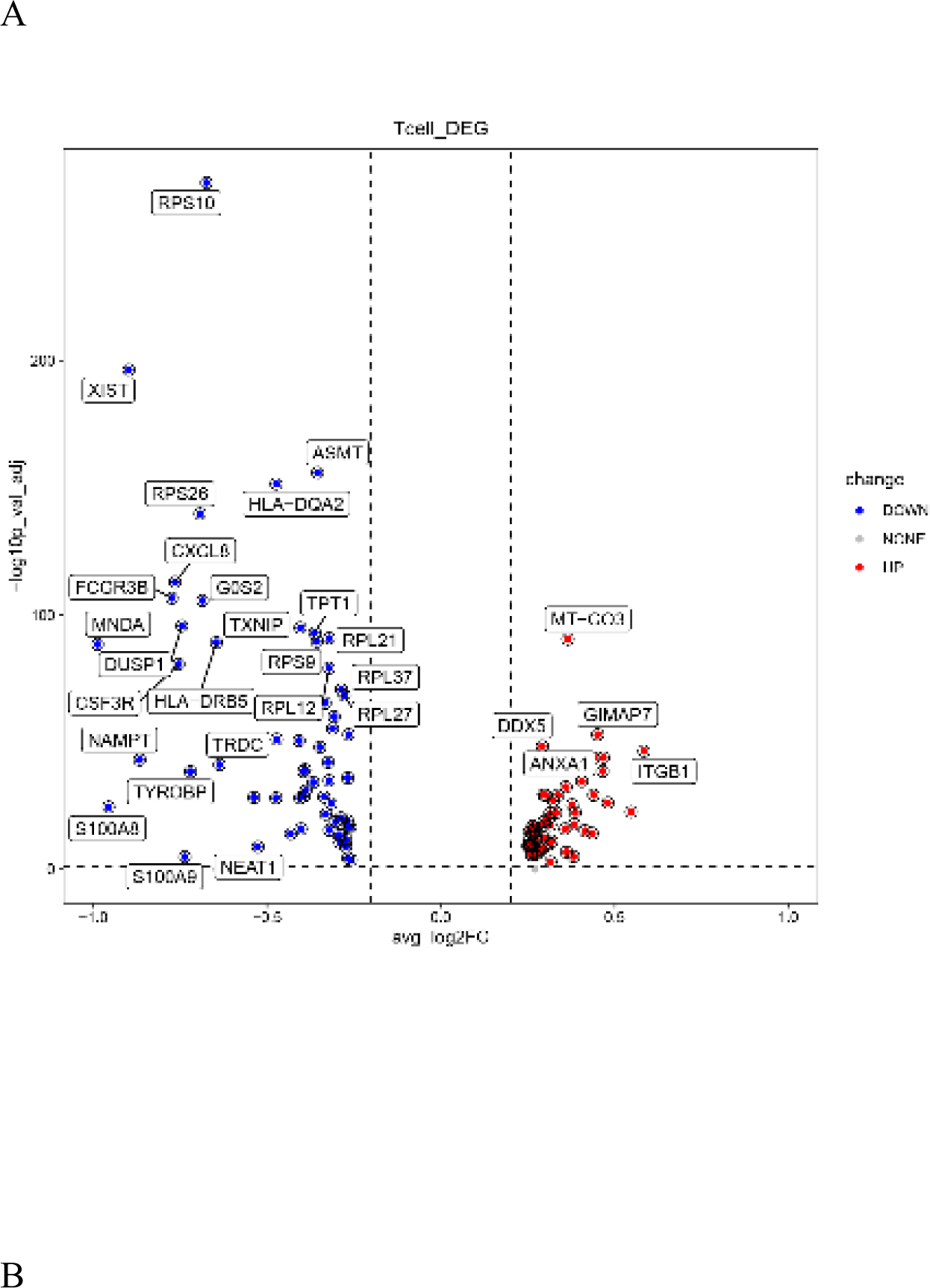
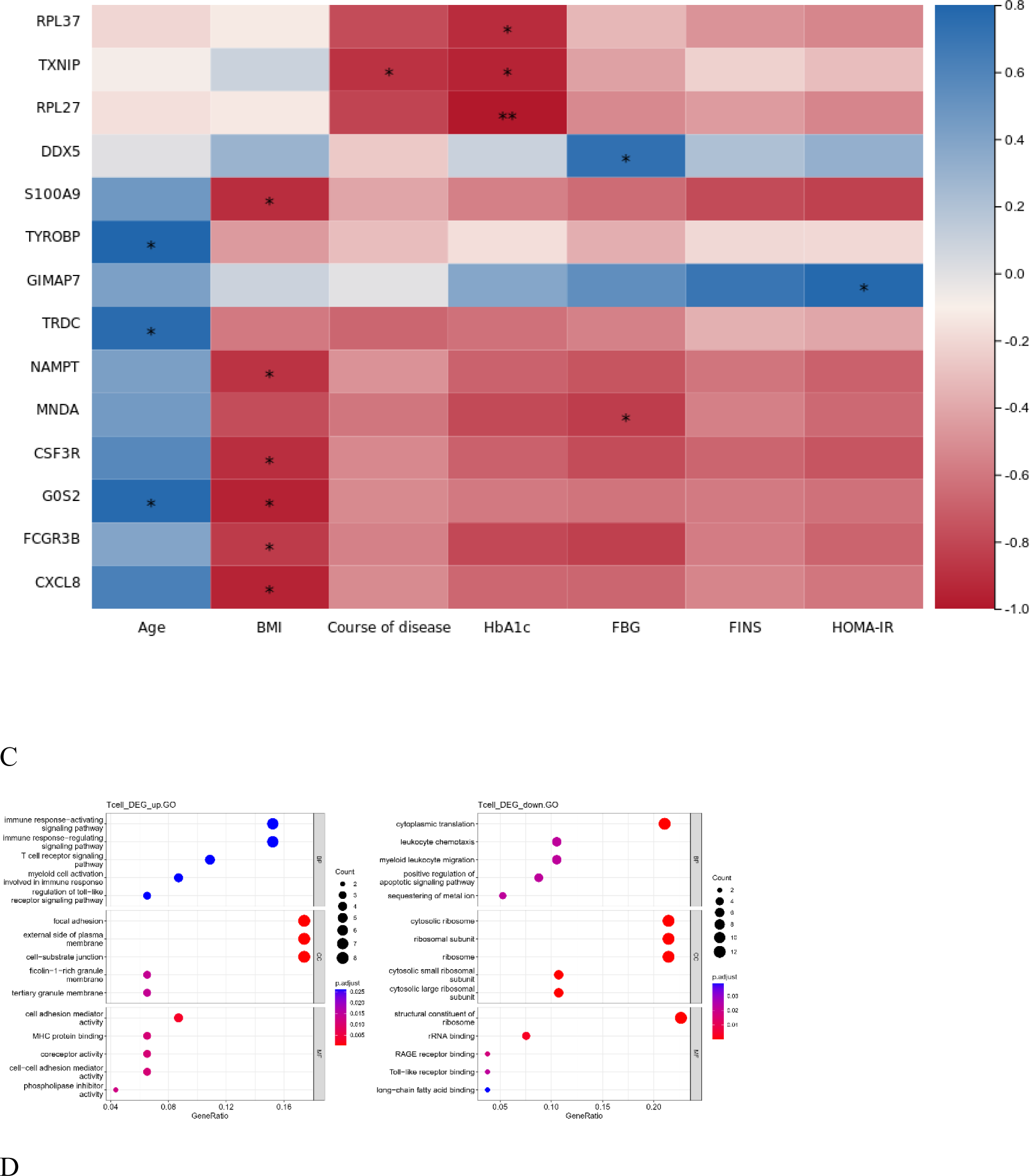
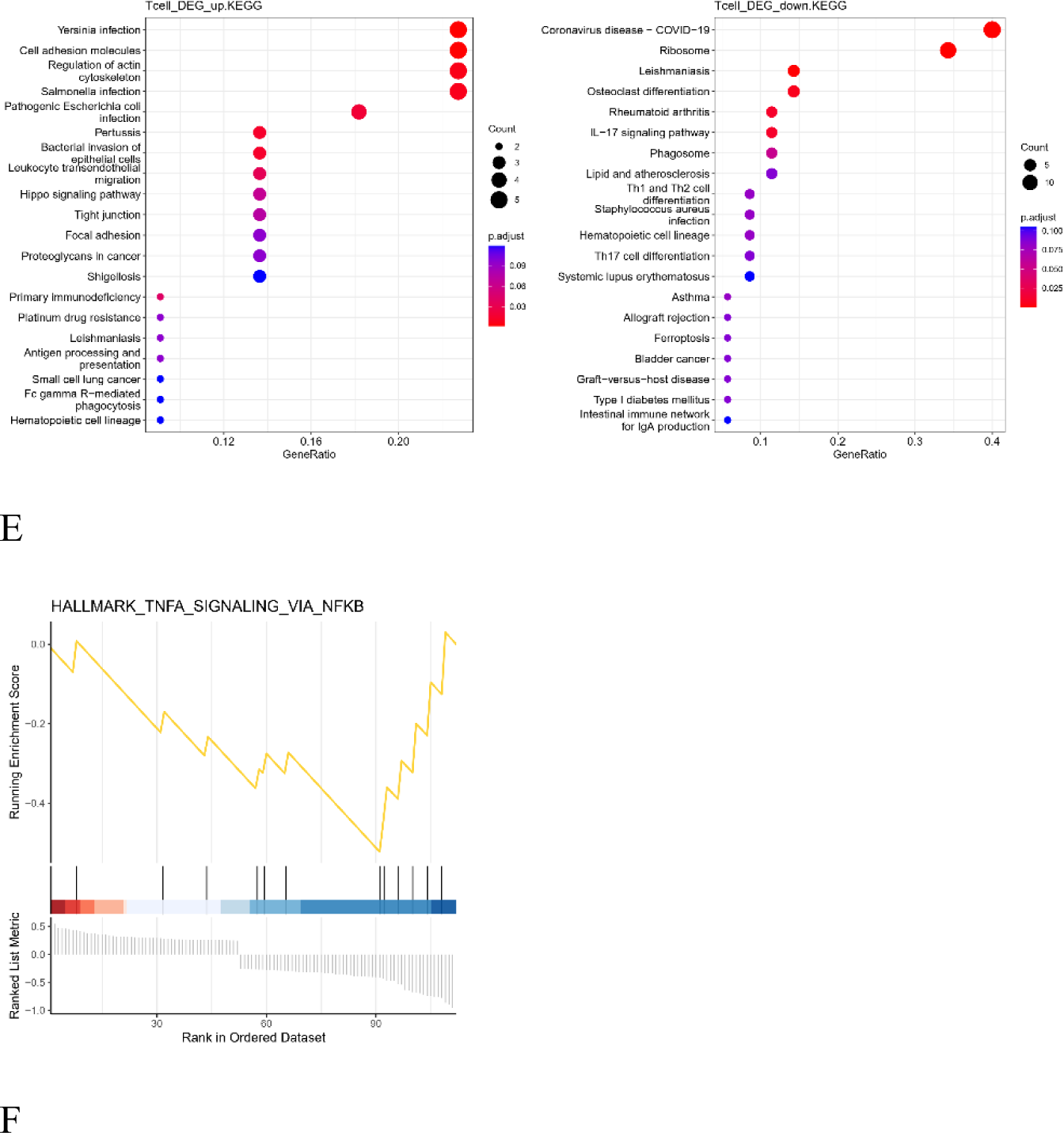
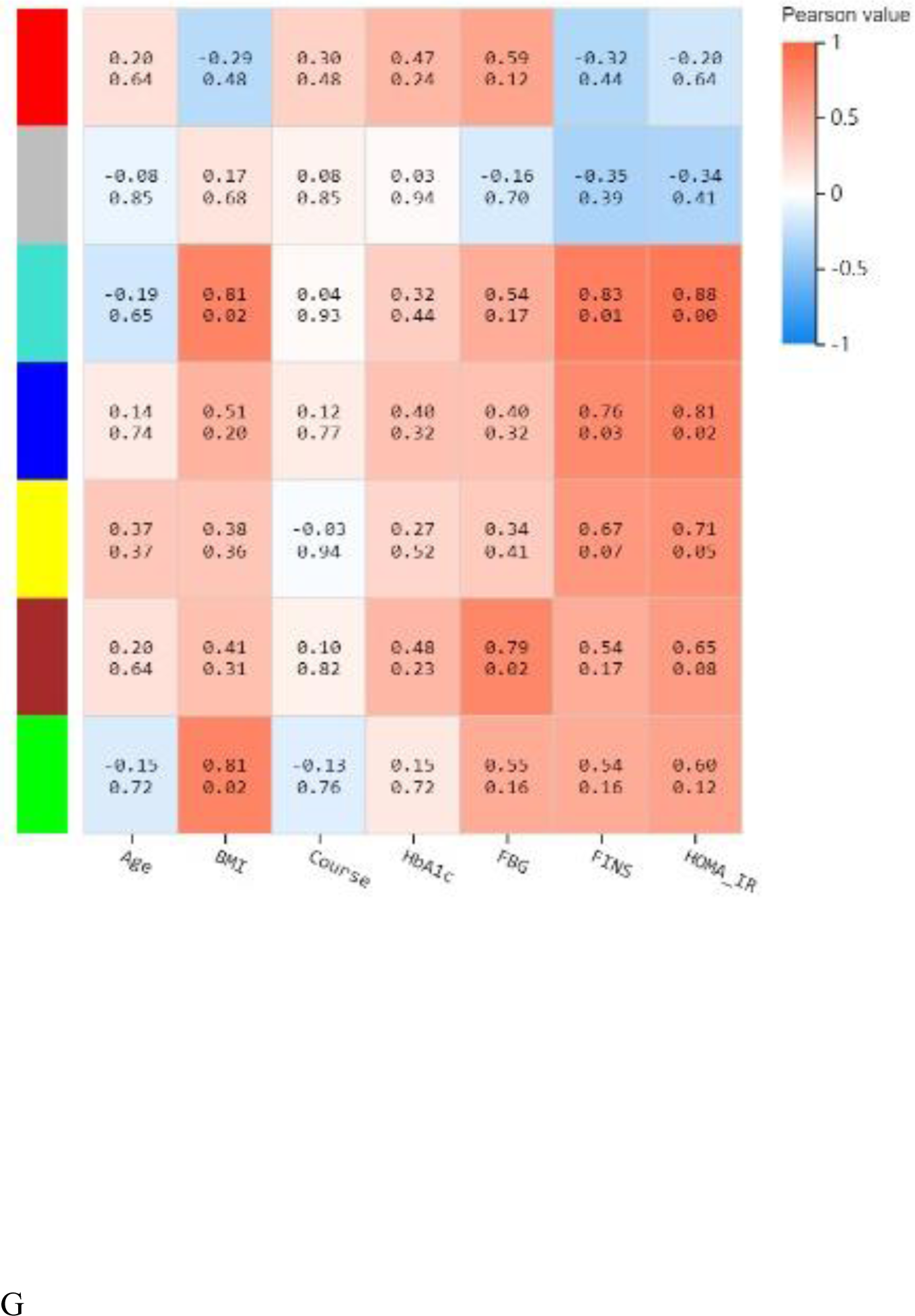
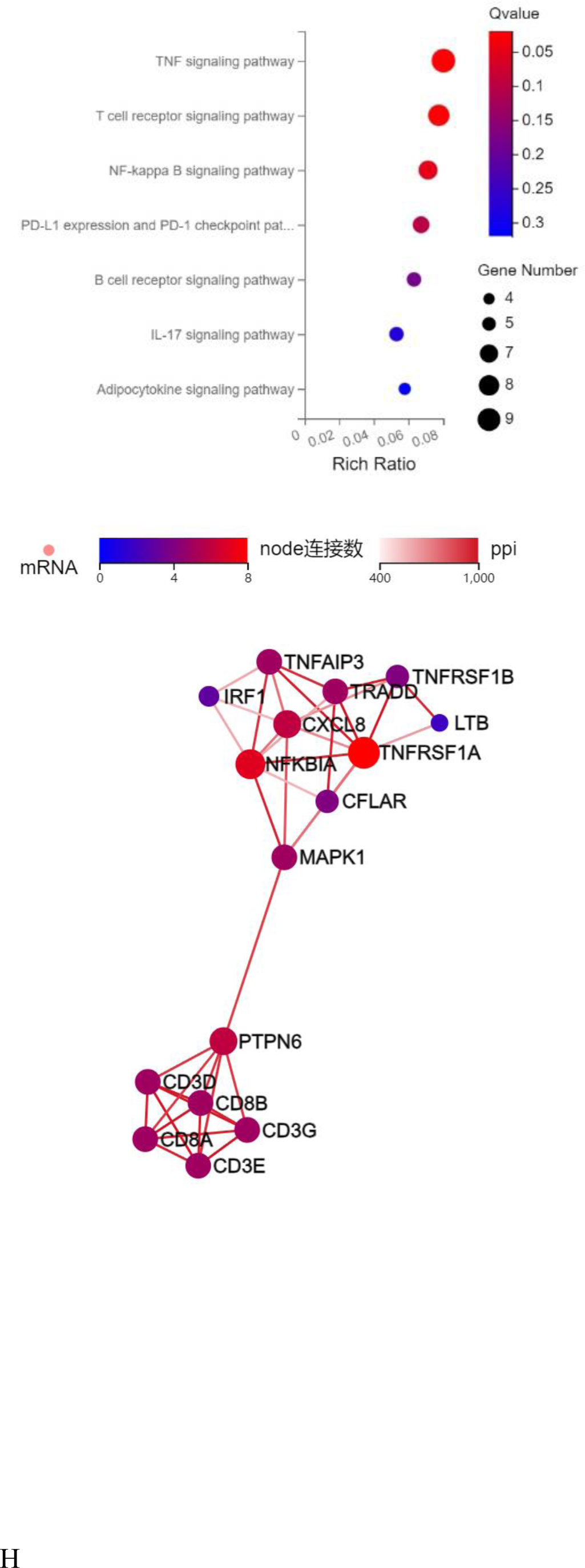
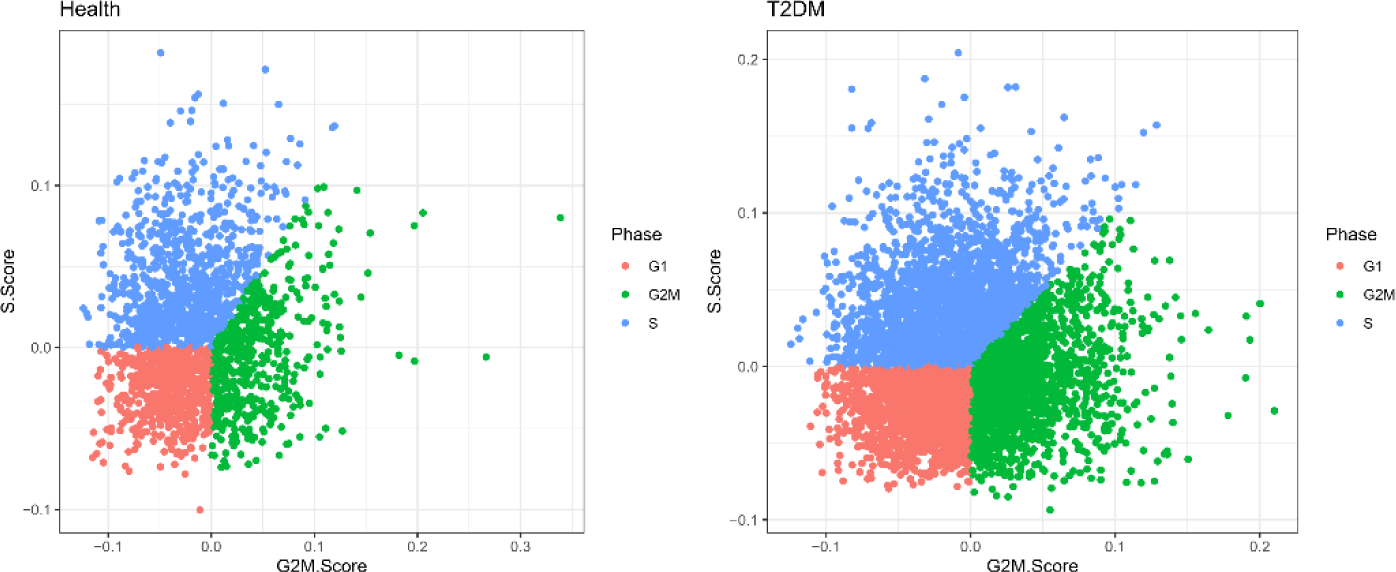
Integrated analysis of T cells. A: The volcano plot shows the differently expressed genes expressed in T cells. B: Correlation analysis between differentially expressed genes and clinical character. C: GO enrichment analysis of the marker genes with up- and downregulated expression in T-cells. D: KEGG enrichment analysis of the marker genes with up- and downregulated expression in T-cells. E: GSEA enrichment analysis of differentially expressed genes. F: Correlation analysis between clinical character and each module. G: The KEGG pathways in module turquoise. D: The cell cycle distribution of T cells.

With respect to T cells, we used known functional genes to identify the CD4+ and CD8+ T-cell populations, including naïve, memory, effector, and Treg populations (Fig. 3D). Moreover, the genes were analysed in CD4+ and CD8+ T cells, and the transcription signal score was calculated. The results suggest that CD4+ T cells in T2DM patients are in memory and naïve states, while CD8+ T cells are in effector and memory states (Fig. 3E). With respect to CD4+ and CD8+ T cells, we screened several specific genes, and their expression is shown in Fig. 3F.

CD4-C5-CCR7-expressing cells are naïve cells, and CD4-C0-AQP3-expressing cells are memory cells. According to their biological background, CD4-C5-CCR7-expressing cells were differentiated into CD4-C0-AQP3-expressing cells. The potential developmental trajectory between healthy controls and T2DM patients is shown in Fig. 5A. The 50 genes whose expression varied with developmental time were grouped into three clusters. These genes were associated with cytoplasmic translation, cell‒cell adhesion mediated by integrin, and regulation of the inflammatory response (Fig. 5B). The figure shows the dynamic expression of six CD4+ T-cell marker genes, and different expression levels can be observed at the seven stages of disease progression. The expression of the RPL32, RPS10, RPS12, RPS14 and RPS23 genes showed an upwards trend, while S100A4 expression showed a downwards trend (Fig. 5C). Since both cell clusters among the CD8+ cells were effector cells, the origin of differentiation could not be determined.

**Figure 5.**
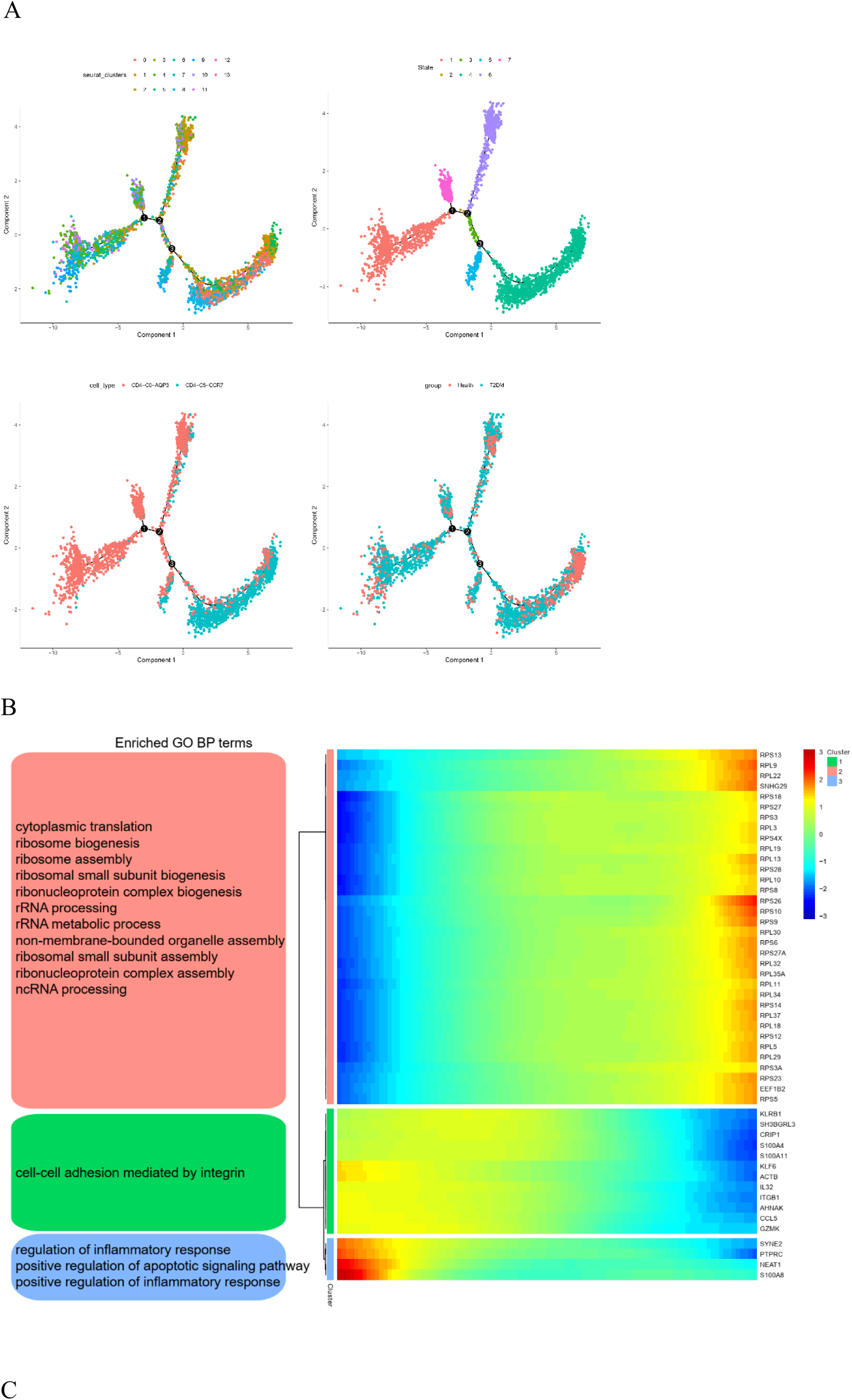
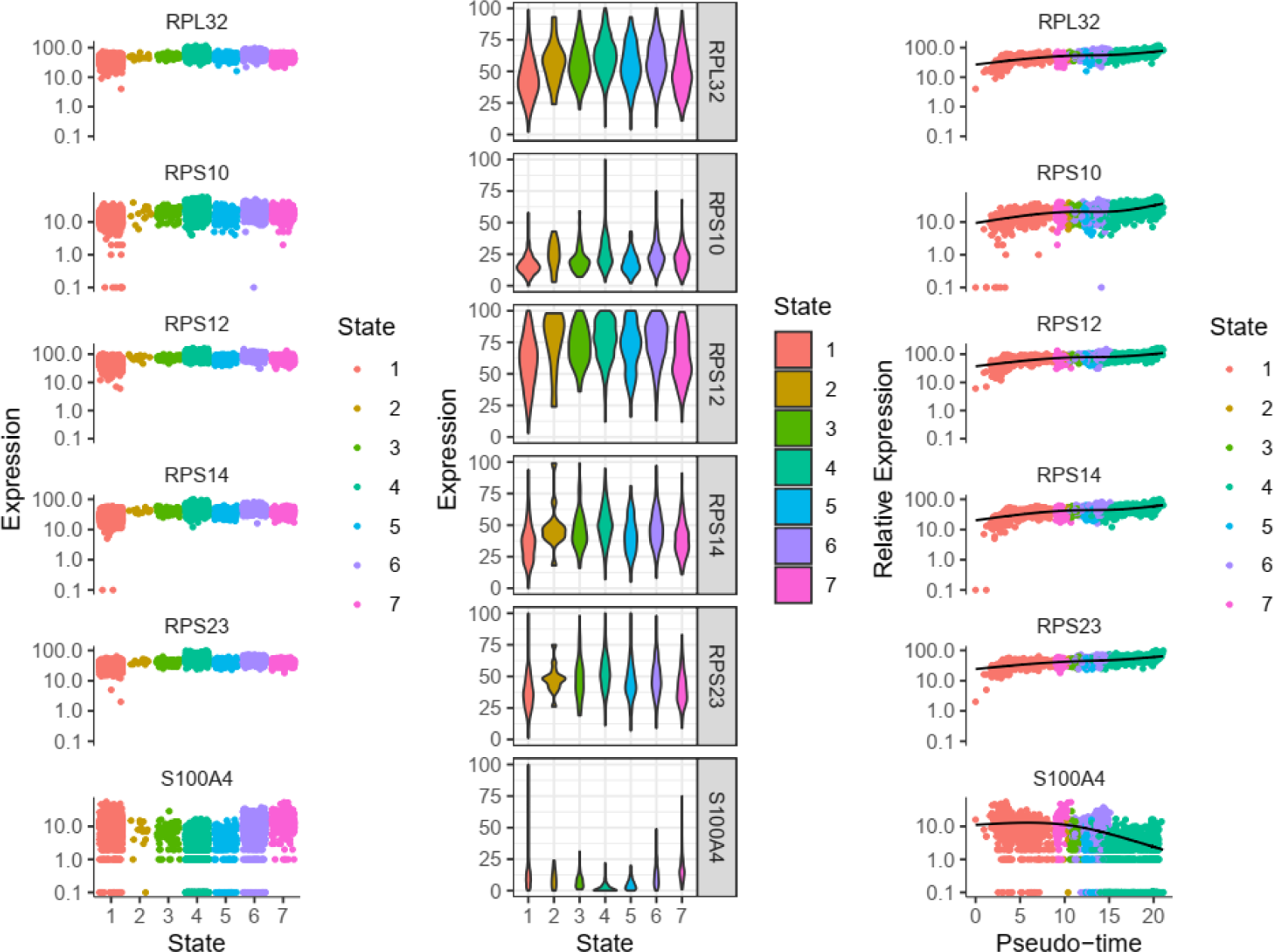
Trajectory inference analysis of CD4+ T cells. A: Trajectory plots showing the different developmental processes of CD4+ T cells. B: Heatmap showing the expression of dynamic genes and the GO analysis results. C: Dynamic expression of the top genes in CD4+ T-cells.

### Clustering and Subtype Analysis of Monocytes/Macrophages

Monocytes/macrophages were the most abundant nonspecific immune cells in our T2DM patient cohort, with a total cell count of 2091. Unsupervised clustering of monocytes/macrophages identified two monocyte clusters (comprising 1918 cells) and 1 macrophage cluster (including 173 cells) (Fig 6A). The expression levels of genes in different cell types are shown by a heatmap. Monocytes/macrophages were annotated separately by canonical genes, and the canonical expression of genes in these cell types was consistent with the annotation (Fig 6B). There were 689 marker genes in monocytes and 70 in macrophages. The bubble diagram shows the average expression levels and percentage of expression of representative marker genes in cells of healthy subjects and T2DM patients (Fig. 6C). With respect to monocytes/macrophages, we used known functional genes to identify monocyte and macrophage populations, including classic monocytes and nonclassical monocytes (Fig. 6D). The results suggest that the CD14+ monocytes in our T2DM cohort were in the classic monocyte state, while the FCGR3A+ monocytes were in the nonclassical monocyte state (Fig. 3E). With respect to monocytes/macrophages, we screened several specific genes, and their expression is shown in Fig. 6F.

**Figure 6.**
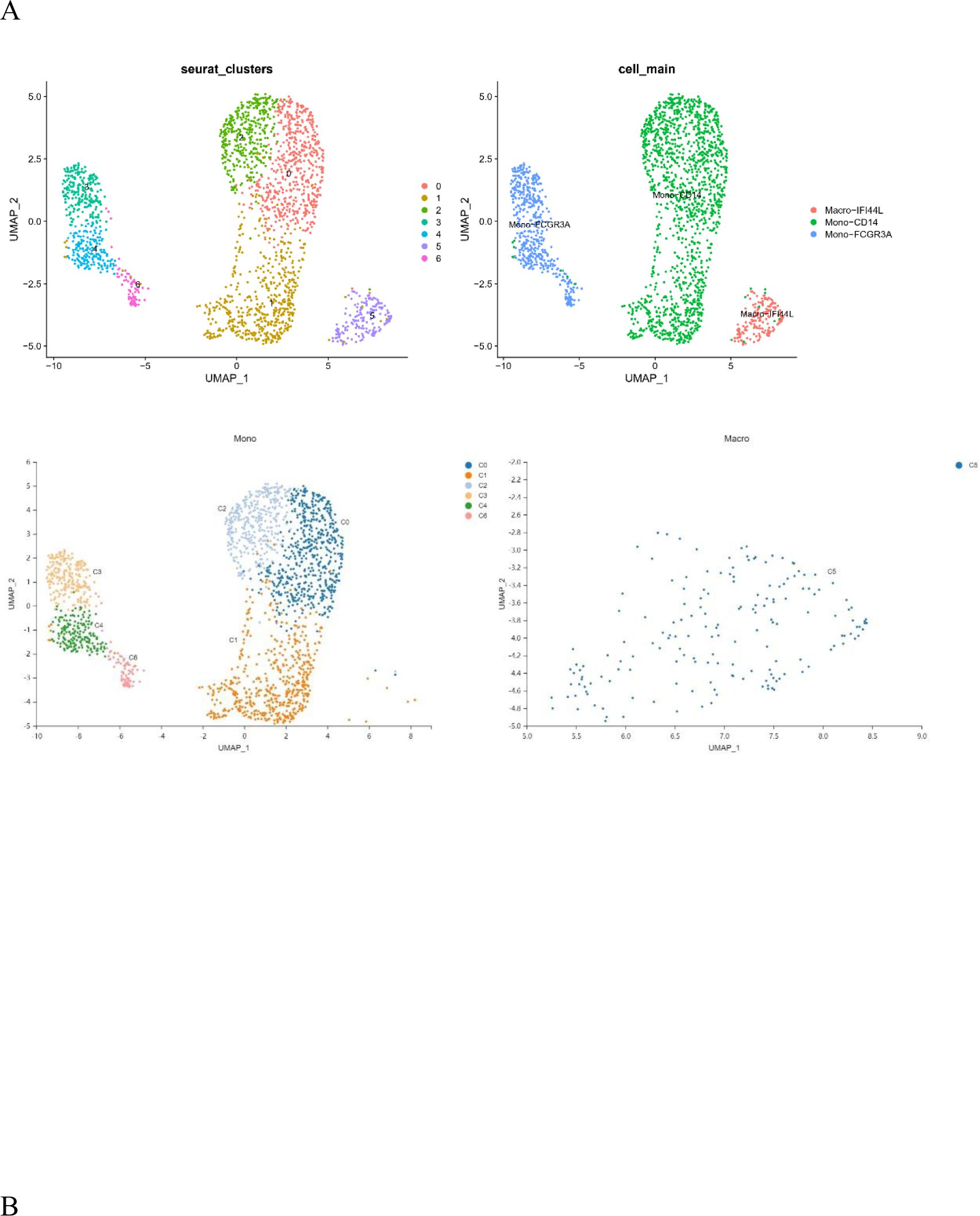
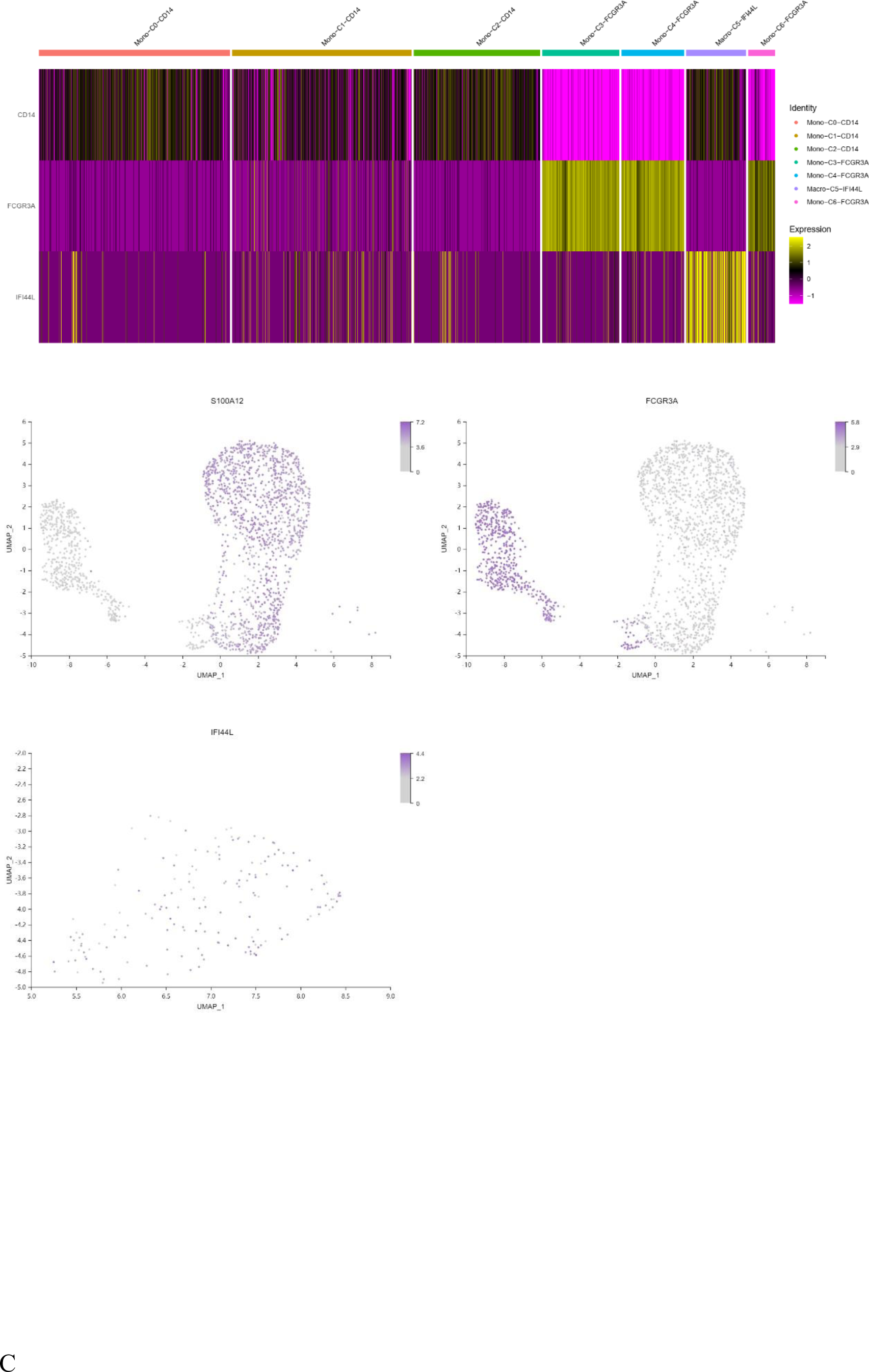
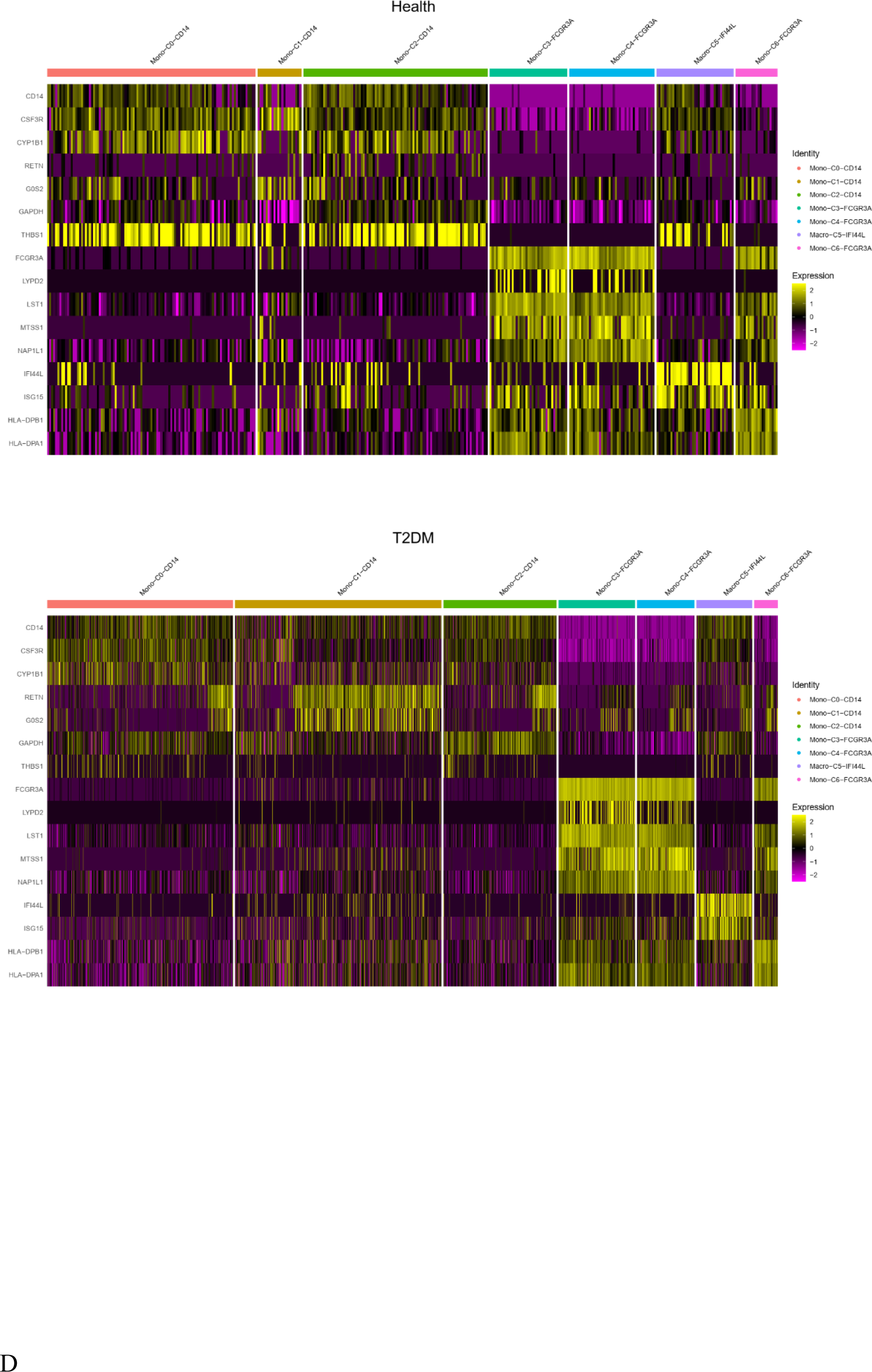
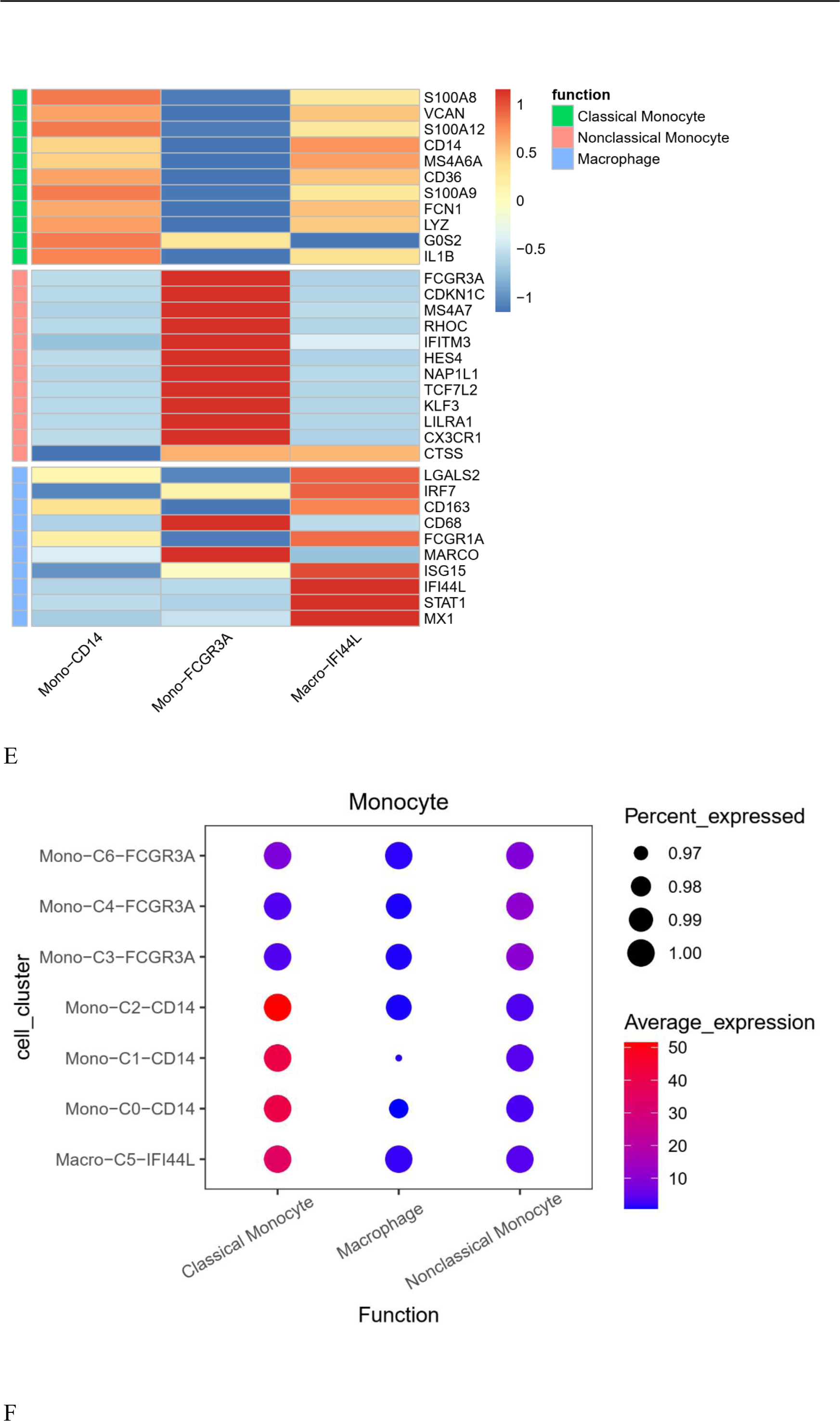
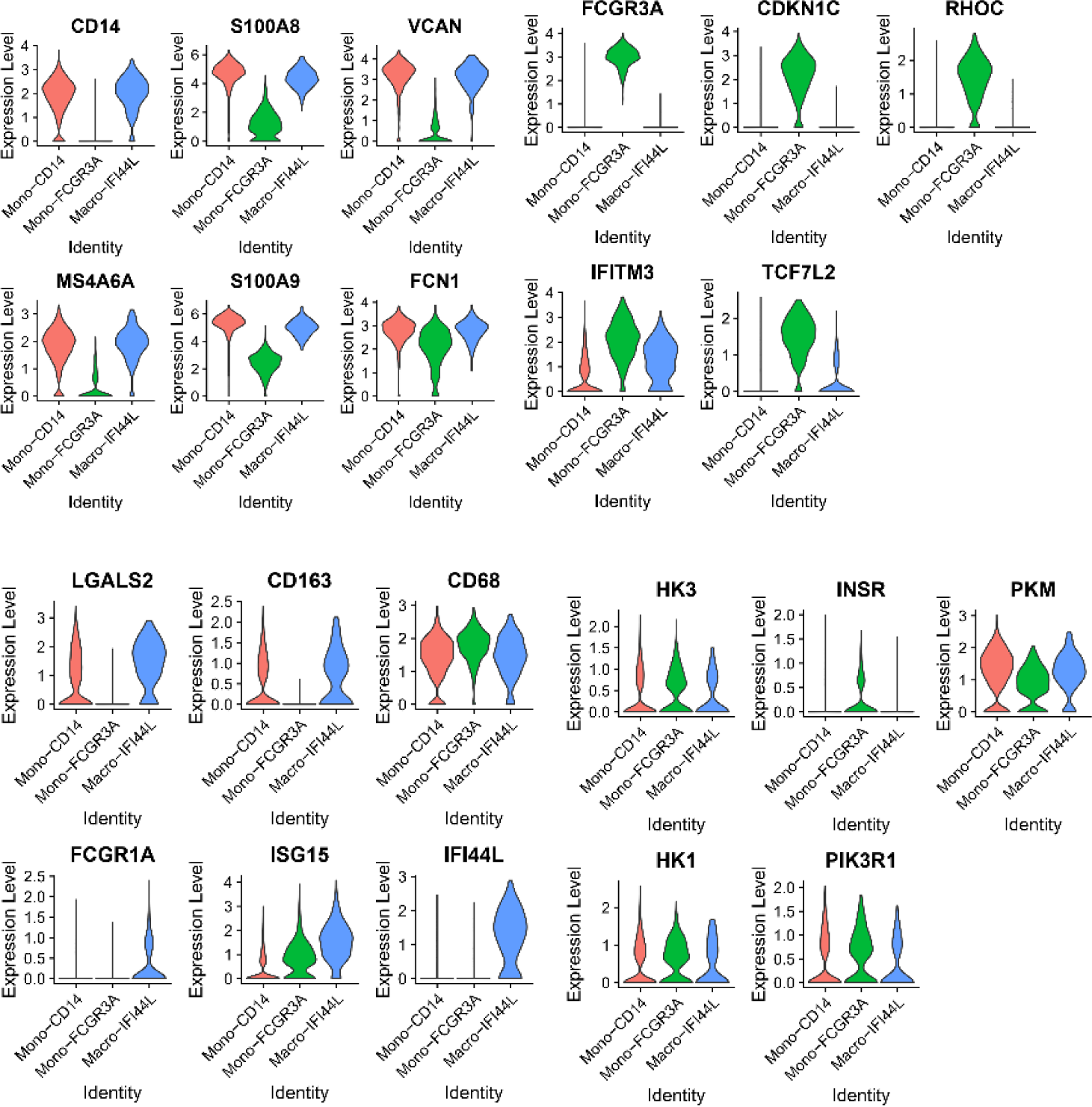
Characterization of monocytes/macrophages. A: UMAP plot of monocytes/macrophages. Two cell clusters were present in monocytes, and one cell cluster was present in macrophages. B: A feature map showing the expression of marker genes for each cell type. The cells were identified by marker genes. C: Dot plot of representative gene signatures in monocyte/macrophage clusters for healthy controls and T2DM patients. D: Heatmap of the expression of selected monocyte/macrophage function-associated genes. E: Dot plot of representative activation stage signatures in monocyte/macrophage clusters. F: Violin plot showing representative genes in monocytes/macrophages.

The differences in monocyte/macrophage gene expression between the T2DM group and healthy control group were compared to construct a volcano plot, as shown in Fig. 7A. Among these genes, 51 had upregulated expression, and 124 had downregulated expression. The CLEC7A, SIGLEC14 and AC018755.4 of genes were negatively correlated with HbA1c. VSTM1 was negatively correlated with FINS and HOMA-IR (Fig. 7B). The GO enrichment results showed that BP was related to granulocyte chemotaxis, cell chemotaxis, leukocyte chemotaxis and viral process. CCs were mainly associated with cytosolic ribosomes, cell−substrate junctions, focal adhesion and tertiary granules. The MFs were specifically related to the structural constituents of ribosomes, enzyme inhibitor activity, S100 protein binding, cytokine activity and cytokine binding cell‒cell adhesion mediator activity (Fig. 7C). The KEGG results showed that the marker genes were enriched in ribosomes, the NOD-like receptor signalling pathway, cytokine‒cytokine receptor interaction, Th17 cell differentiation, the TNF signalling pathway, and Th1 and Th2 cell differentiation (Fig. 7D). The first gene set, HALLMARK_INTERFERON_GAMMA_RESPONSE, was utilized to generate a GSEA graph via HALLMARK pathway enrichment analysis (Fig. 7E). The results of WGCNA showed that genes were divided into 7 modules. FBG was significantly negatively correlated with module red. FINS and HOMA-IR were significantly negatively correlated with module brown, positive correlated with module turquoise (Fig. 7F). The KEGG result of genes in the module red is involved in the Chemokine signaling pathway (Fig. 7G). We further analysed the cell cycle progression of monocytes/macrophages in healthy controls and T2DM patients. In T2DM, monocytes/macrophages remained in the S and G2M stages (Fig. 7H).

**Figure 7:**
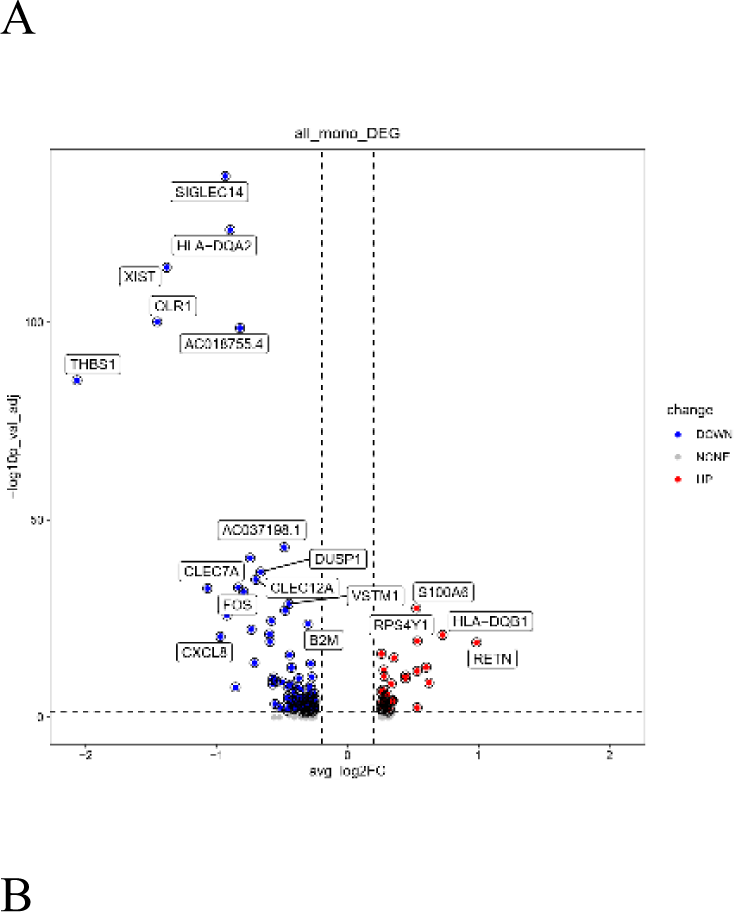
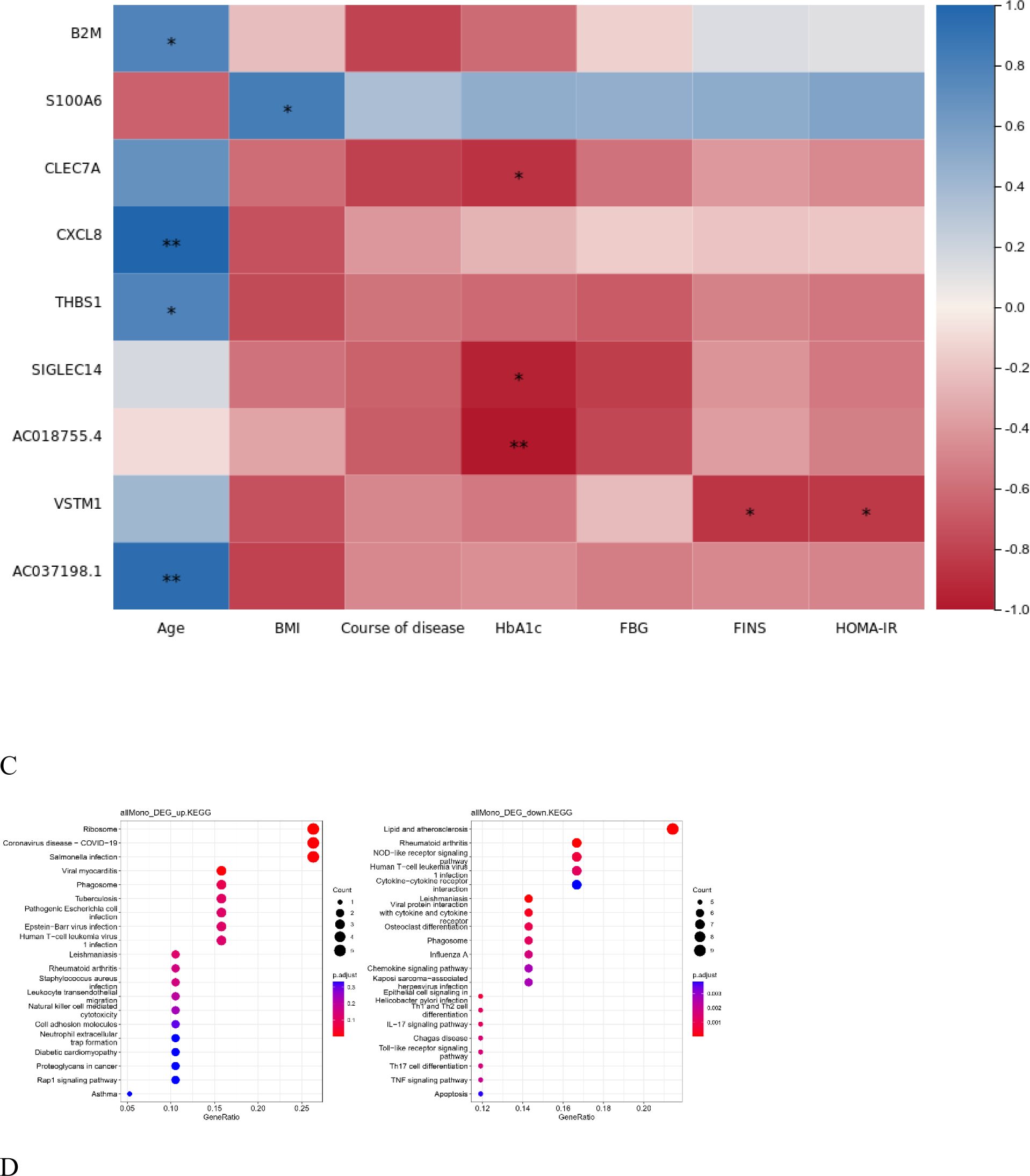
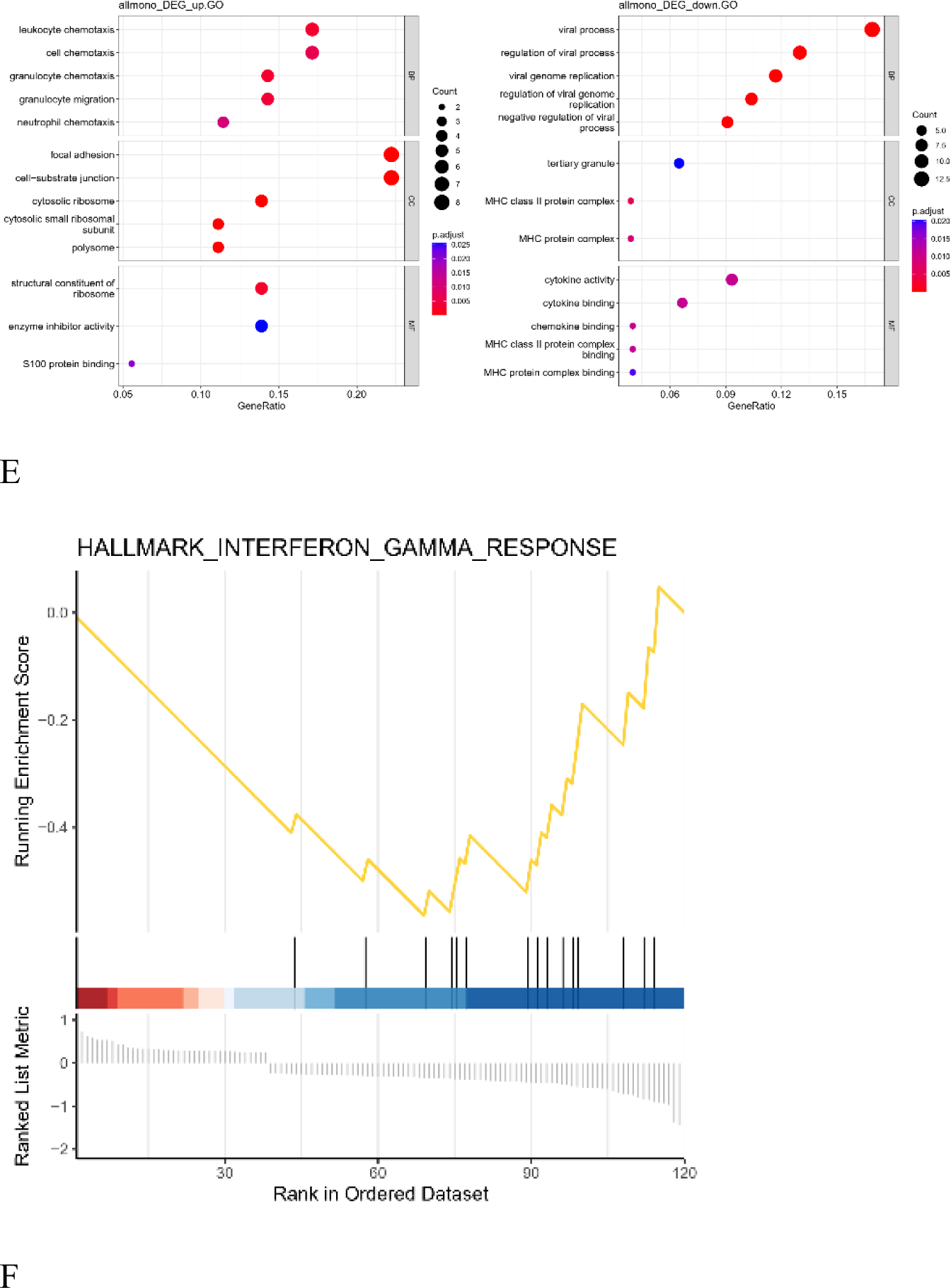
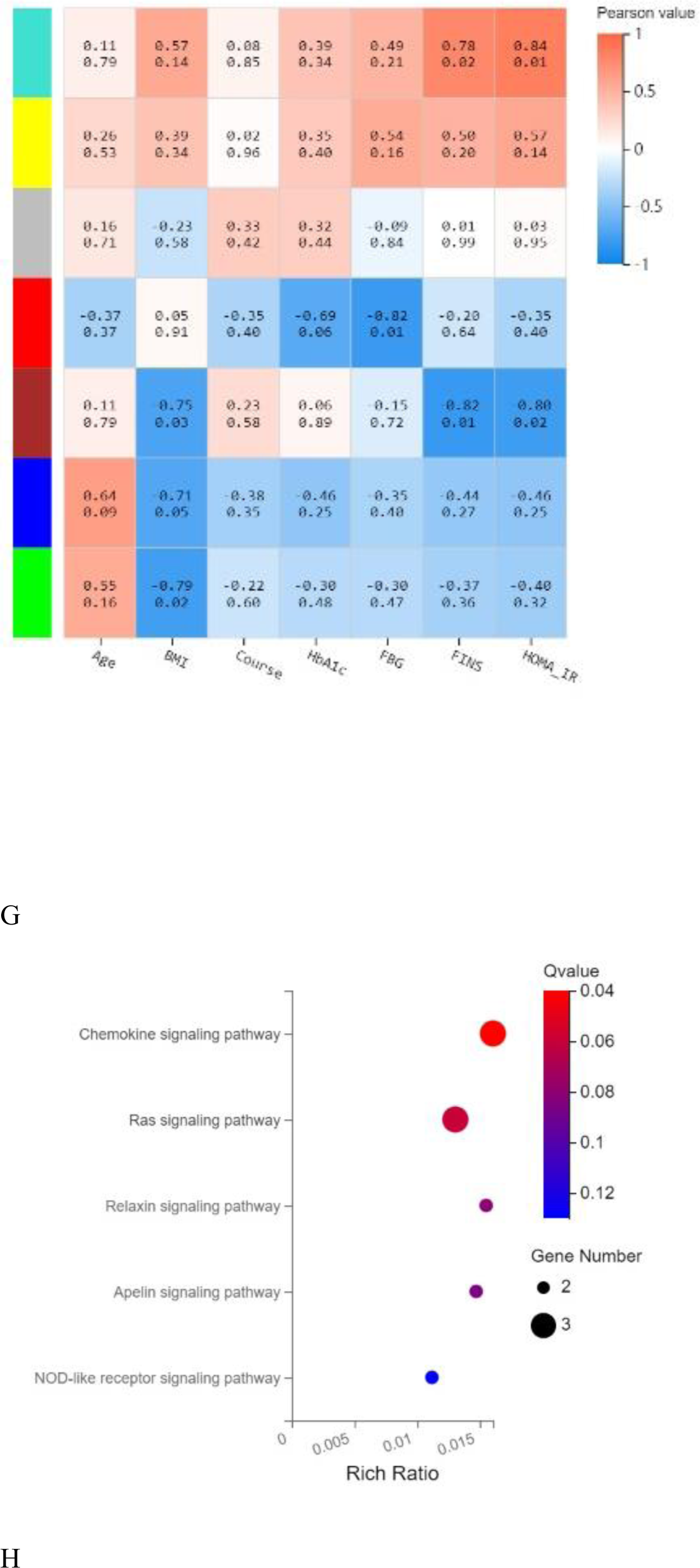
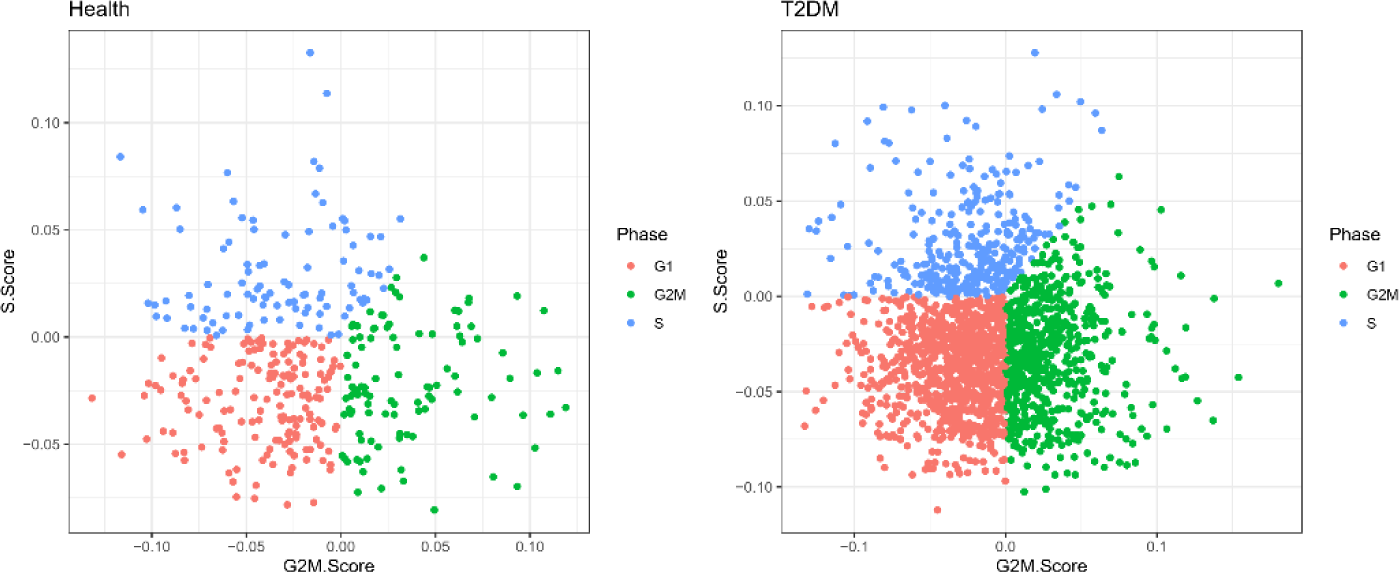
Integrated analysis of monocytes/macrophages. A: The volcano plot shows the differentially expressed genes in monocytes/macrophages. B: Correlation analysis between differentially expressed genes and clinical character. C: GO enrichment analysis of the marker genes with up- and downregulated expression in monocytes/macrophages. D: KEGG enrichment analysis of marker genes with up- and downregulated expression in monocytes/macrophages. E: GSEA enrichment analysis of differentially expressed genes. F: Correlation analysis between clinical character and each module. G: The KEGG pathways in module red. H: The cell cycle phase of monocytes/macrophages.

We constructed quasitime differentiation and developmental trajectories of monocytes/macrophages. According to the biological background analysis combined with this trajectory, there was partial differentiation from CD14+ monocytes and FCGR3A+ monocytes to IFI44L+ macrophages. Because the sample used was peripheral blood, the number of macrophages present was very small. The potential developmental trajectory between healthy controls and T2DM patients is also shown (Fig. 8A). The 50 genes that varied with developmental time were grouped into four clusters. These genes are associated with the negative regulation of hydrolase activity, neutrophil migration, polyamine biosynthetic processes, and positive regulation of cytokine production (Fig. 8B). The figure shows the dynamic expression of six monocyte/macrophage marker genes, and the expression levels are different at various differentiation stages. Fig. 8C shows the dynamic expression of six monocyte/macrophage marker genes, which differed among the three stages. The expression of the C1QA, HES4 and RHOC genes showed an upwards trend, while S100A12, S100A8, and S100A9 expression showed a downwards trend.

**Figure 8.**
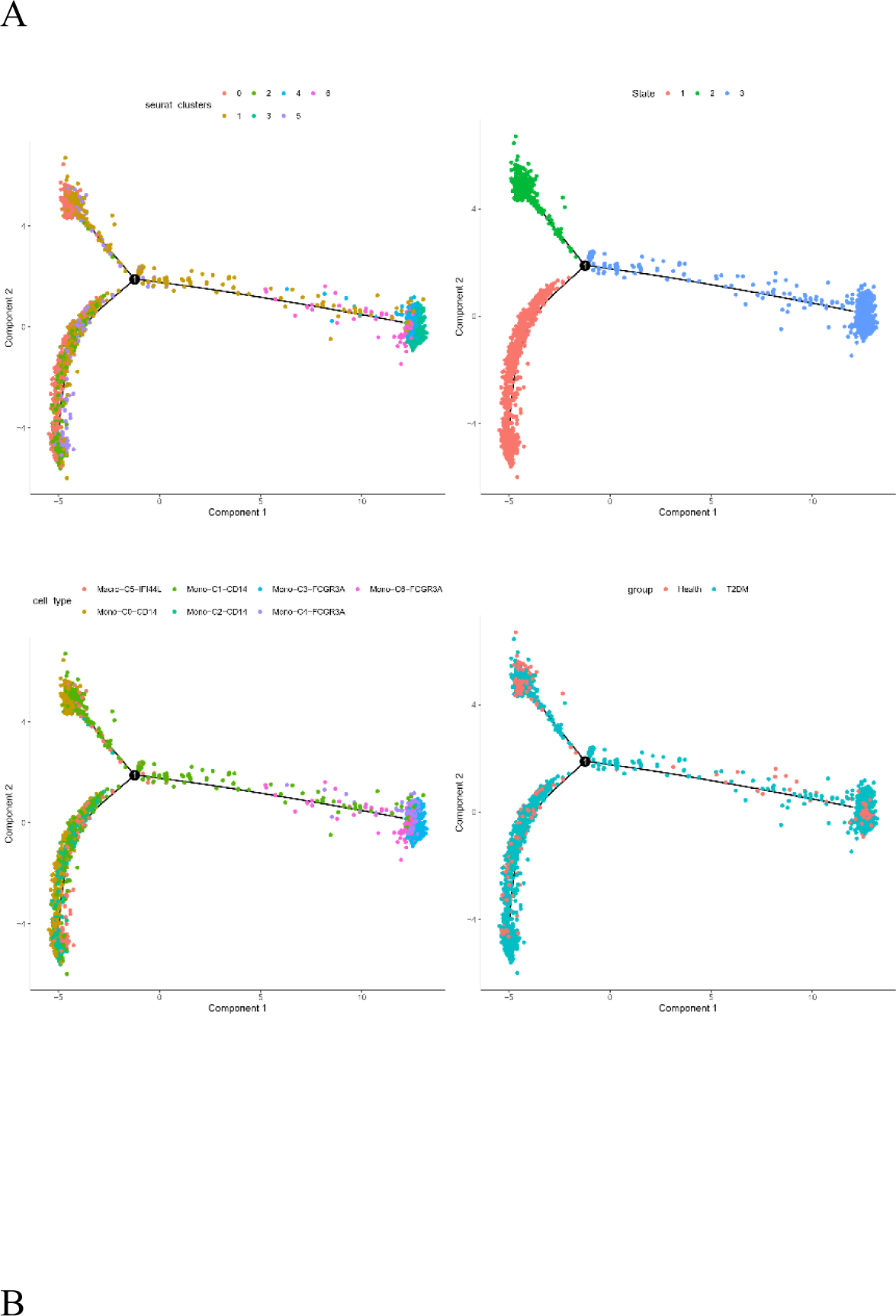
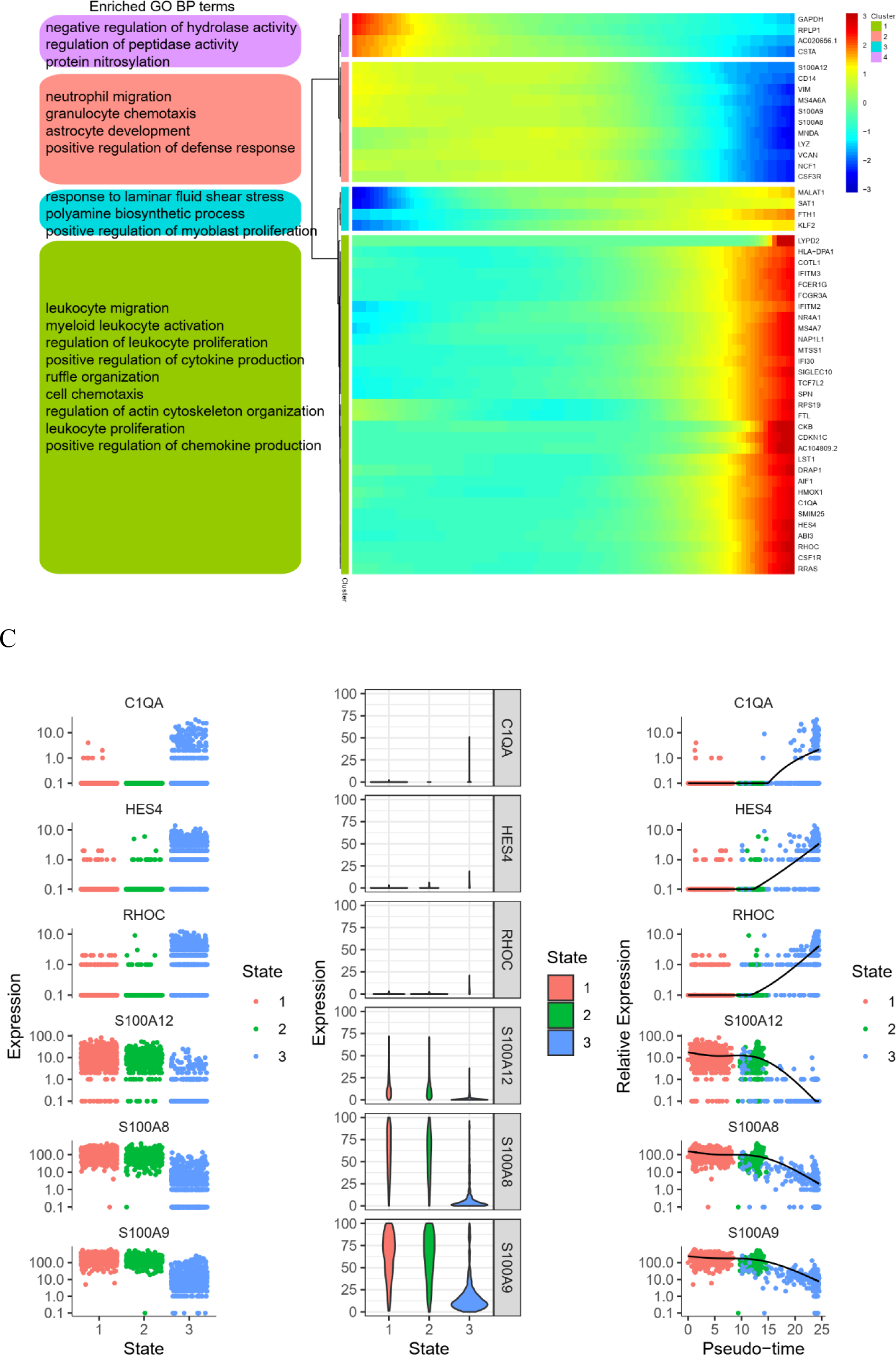
Trajectory inference analysis of monocytes/macrophages. A: Trajectory plots showing different developmental processes in monocytes/macrophages. B: Heatmap showing the expression of dynamic genes and the GO analysis results. C: Dynamic expression of the top genes in monocyte/macrophage.

### Cell‒cell interaction and communication properties

The results revealed the number and weight/strength of cell‒cell interactions between monocytes/macrophages and between CD4+ and CD8+ T cells and NK cells (Fig. 9A and 9B). The interactions were mainly present in the MIF signalling pathway (Fig. 9C). Ligand receptor-mediated interactions were found mainly in MIF-(CD74 + CD44), MIF-(CD74 + CXCR4), ANXA1-FPR1 and LGALS9-CD45 cells (Fig. 9D). Among them, the ligand‒receptor pair MIF-(CD74 + CD44) contributed the most to cell‒cell interactions. The gene expression of MIF, CD74, CXCR4, CD44 and CXCR2 was represented in the MIF signalling pathway (Fig. 9E). CD4+ and CD8+ T cells were the major transmitters of the MIF signalling pathway, and the major receivers, mediators and influencers were monocytes/macrophages (Fig. 9F).

**Figure 9.**
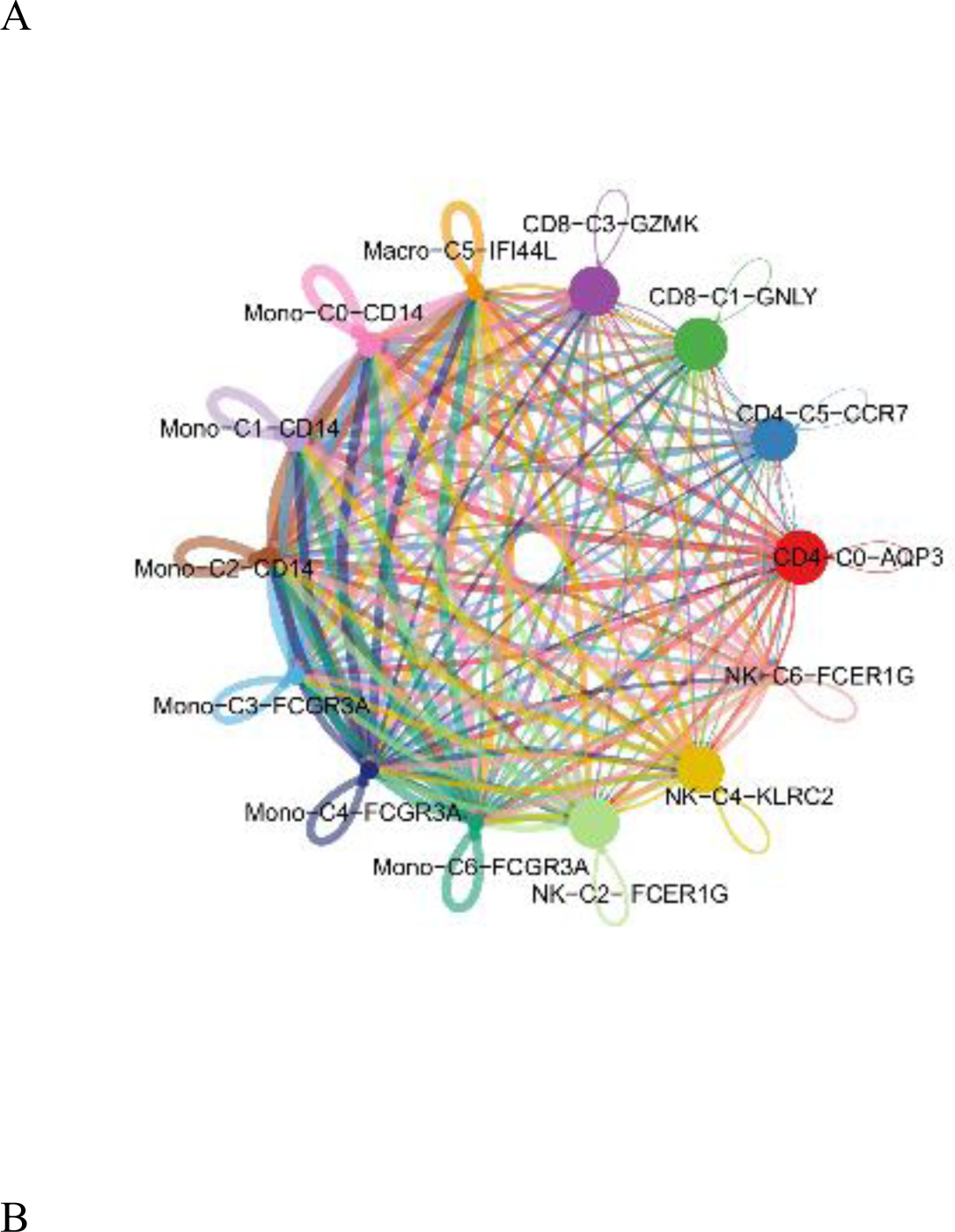
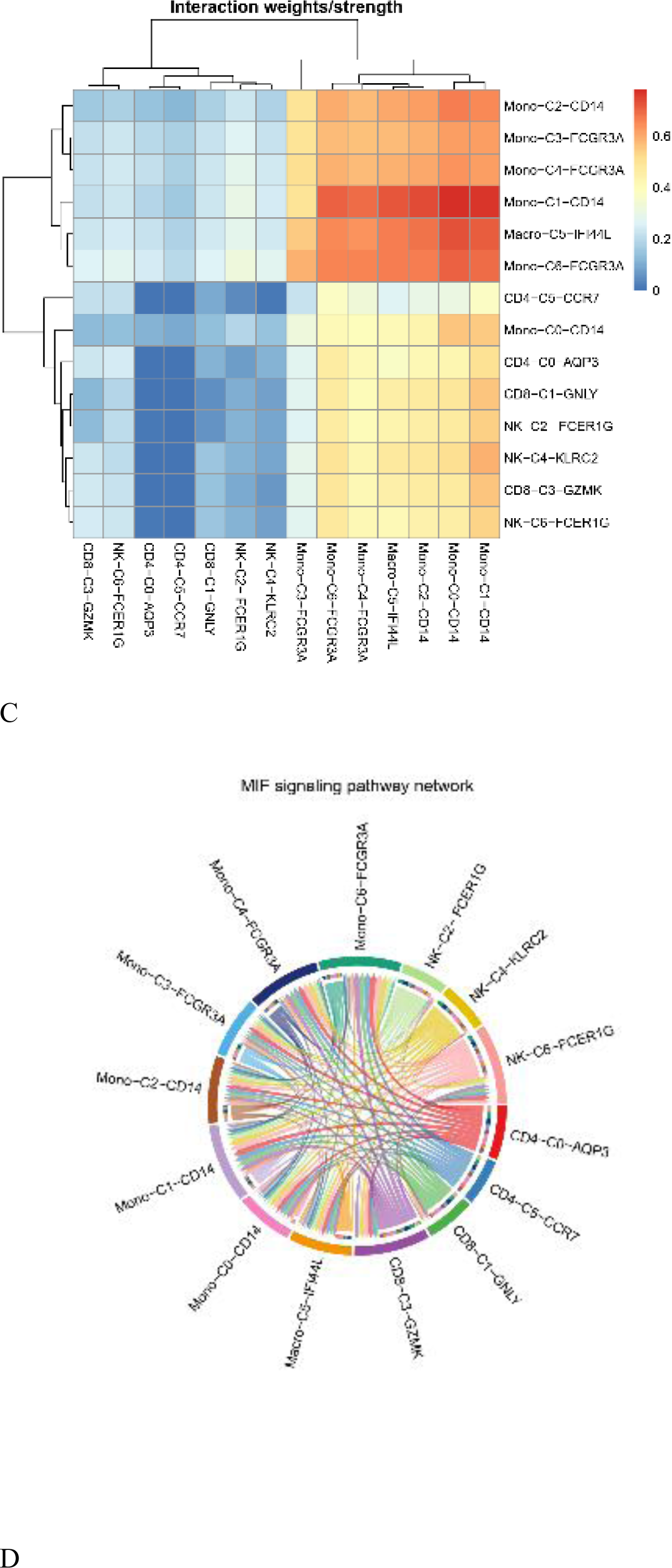
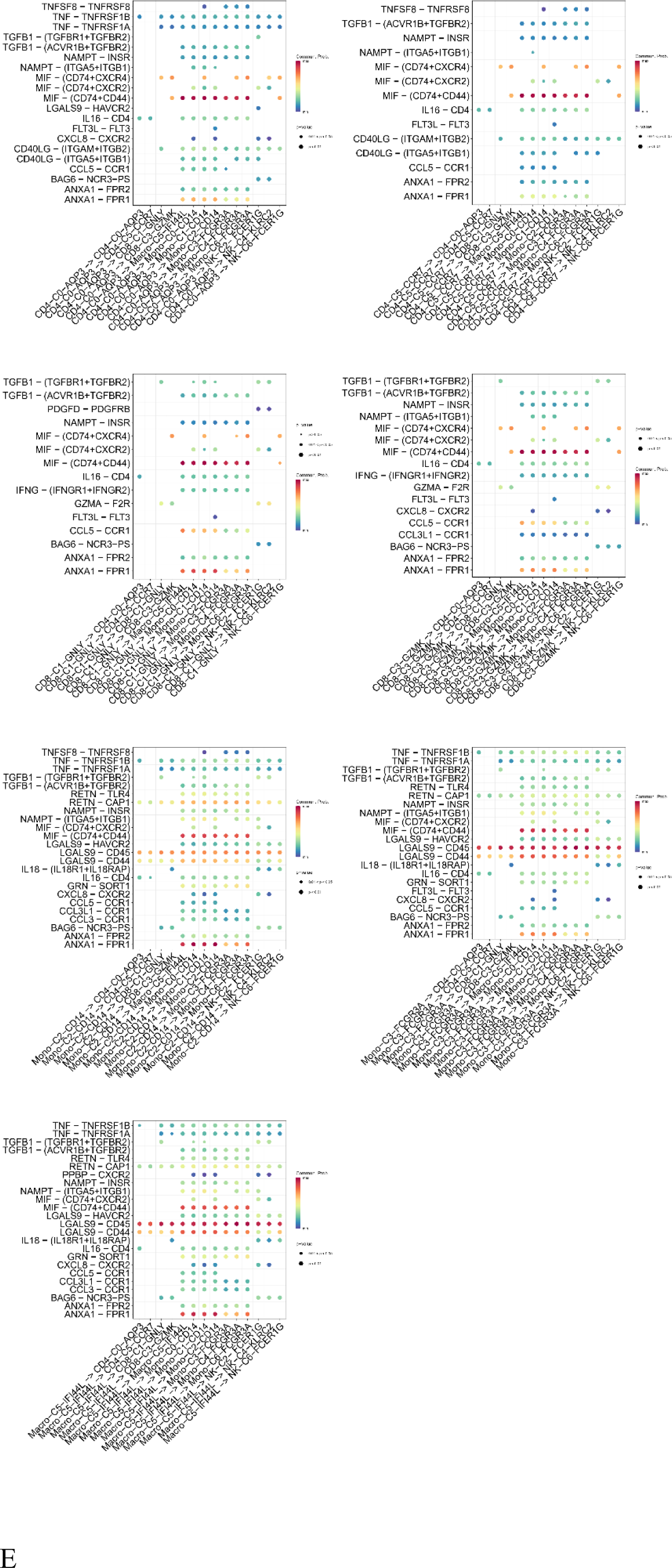
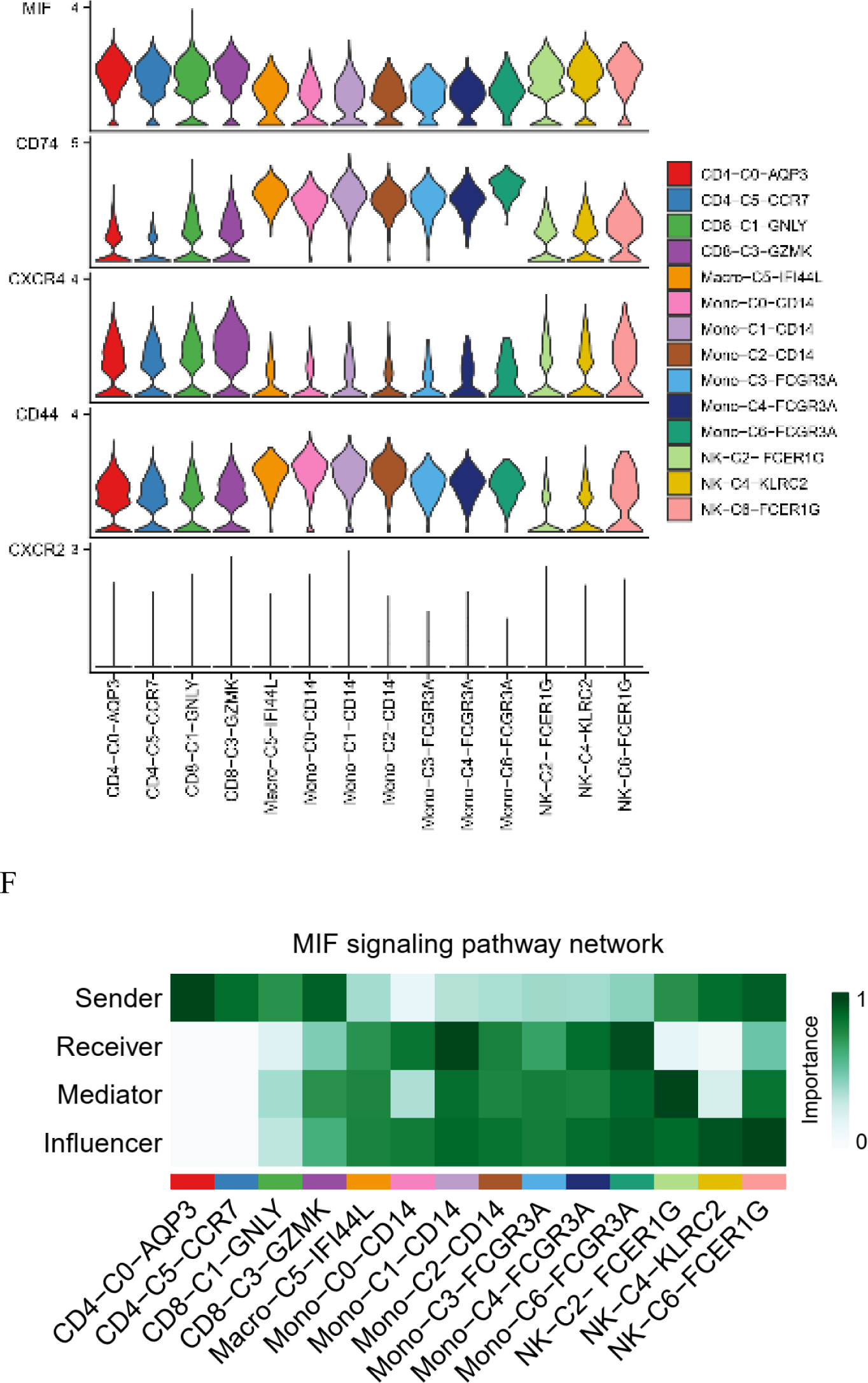
Cell‒cell communication analysis. A: The number of interactions. B: The weights/strengths of interactions. C: Interactions in the MIF signalling pathway. D: The relationships between ligand‒receptor pair–mediated interactions and cells. E: The expression of genes in the MIF signalling pathway. F: The role of cells in the MIF signalling pathway.

## Discussion

Compared with type 1 diabetes, T2DM has a higher incidence and longer duration, and the complications tend to be worse. At present, scRNA-seq is widely used to characterize the basic properties of cells, and the regulation of islet cells in diabetes by sub cohorts of cells has been reported [17–18]. Since islet cells are difficult to obtain from the human body, systematically elucidating the regulation of peripheral blood mononuclear cells in patients with T2DM is important. Previous evidence suggests that the pathogenesis of T2DM is related to the immune system [19–20].

Single-cell clustering analysis revealed that 14 cell clusters were annotated to 7 different cell types, namely, neutrophils, T cells, NK cells, monocytes/macrophages, B cells, cDC1 cells and platelets. Interestingly, we found that monocytes/macrophages and T cells were expressed at higher levels in T2DM patients than in healthy controls.

Based on the expression levels of 339 genes in the endocrine and metabolic disease systems, KEGG enrichment analysis demonstrated that these genes are more involved in type I diabetes mellitus, the insulin signalling pathway, the AGE/RAGE signalling pathway in diabetic complications, insulin resistance, T2DM, maturity-onset diabetes of the young, etc. These diseases and signalling pathways are directly or indirectly related to T2DM. We screened 11 genes related to T2DM. Among the Type 2 Diabetes Knowledge Portal-predicted effector genes, INSR and IRS2 are strongly involved, whereas MAPK3, PIK3R1, HK1, PIK3CD and TNF are moderately involved [21]. In addition, the relationship between SOCS3 and HbA1c was very strong. Based on the 11 genes, we constructed a KEGG pathway network. In addition to their direct association with T2DM, 9 genes are related to the insulin signalling pathway. Insulin can bind to its receptor on the cell surface (InsR) and undergo a series of signalling cascades to lower blood glucose levels. For example, insulin inhibits the FoxO signalling pathway and reduces gluconeogenesis activity [22]. Metformin via the activation of FoxO signalling pathway for regulating blood glucose and protecting microvascular endothelial cells [23]. IRS2 is an insulin substrate that regulates blood glucose levels. Mice with IRS2 knockout exhibited insulin resistance [24–25]. The expression of IRS2 gene was up-regulated in T2DM patients after taking metformin orally [26]. HIF-1 regulates target genes involved in inflammation, and notably, increased HIF-1 signalling promotes the development of metabolic diseases [27], especially in the livers of T2DM patients [28]. Metformin can reduce HIF-1α gene expression[29].

Both the innate immune response and adaptive immunity are involved in inflammation. Innate immunity may cause inflammation via endogenous danger signals. Adaptive immunity also provokes inflammation via cytotoxicity, cytokines and other mediators [30]. There is growing evidence supporting the idea that T2DM is a chronic inflammatory disease that results in insulin resistance and hyperglycaemia [31]. Recently, scRNA-seq has revealed that the populations of CD4+ T cells and CD8+ T cells are expanded in T2DM patients [32–33]. In this study, the activation stages of T cells included naïve, memory, and effector T cells. CD4+ effector T cells are the main cells that exert direct immune effects. Once activated, CD4+ effector T cells and Th1 cells exhibit many significant signs and responses to immune inflammation [34]. Compared with non-T2DM patients, T2DM patients had elevated percentages of CD4+ effector T cells [35]. When stimulated by antigens, memory CD4+ T cells in peripheral blood produce effector cytokines for immune protection. A high number of memory CD4+ T cells was associated with a decreased risk of diabetes [36]. Tregs play a protective role against insulin resistance in the pathogenesis of T2DM [37]. The accumulation of cytotoxic CD8+ effector T cells induces inflammation and insulin resistance [38].

Monocyte and macrophage activation might be indicators in T2DM [39]. In an inflammatory state, monocytes are recruited to the tissue, which subsequently promotes an increase in the macrophage population. Therefore, circulating blood monocyte levels can be used as a parameter to reflect the activation of tissue immunity [40]. Macrophages are the major inflammatory cell type in adipose tissue and the liver. For example, the number of macrophages in adipose tissue increased approximately 45% in T2DM patients [41]. Liver inflammation presumably occurs during the early stages of metabolic dysfunction [42].

There were lineage connections within monocyte/macrophage and T-cell populations. Some of the genes activated in CD4+ cells overlapped with genes in CD8+ cells or those that were activated in monocytes and overlapped with genes in macrophages. The enriched genes included regulatory molecules and diabetes-related receptors, such as IL7R, LTB, ITGB1, PKM, FCN1, HK3, HK1 and PIK3R1. The expression of PIK3R1 can regulate the activity of PI3K signalling pathways and further affect glucose metabolism [43]. In contrast, CD4+ cells expressed markedly higher levels of CD4, CCR7, SELL, LEF1, TSHZ2, MAL, TCF and AQP3 than did CD8+ cells, while CD8+ cells expressed greater amounts of GZMK, GNLY, and NKG7 than did CD4+ cells. FCGR3A+ monocytes expressed markedly higher levels of FCGR3A, CDKN1C, RHOC, HES4, TCF7L2 and INSR than did CD14+ monocytes and macrophages, while CD14+ monocytes and macrophages expressed higher amounts of S100A8, S100A9, VCAN, CD14, MS4A6A, LGALS2, and CD163 than did FCGR3A+ monocytes. TCF7L2 is a potent locus for T2DM risk and is activated by the Wnt signalling pathway [44]. Glycolysis/gluconeogenesis is a central pathway in energy metabolism in the body. The glycolysis/gluconeogenesis pathway was activated in T2DM patients compared with healthy controls. The literature has pointed to a link between glycolysis/gluconeogenesis and prediabetes or T2DM [45]. Glycolysis in a hyperglycaemic state is closely related to the function of macrophages [46]. MGST1, which is positively correlated with FBG, has been validated as a key T2DM gene involved in problems with islet cells [47–48]. MGST1 is a membrane-bound transferase that regulates oxidative stress [49] and is closely related to chronic inflammation.

We have screened some candidate genes by the correlation analysis between differentially genes and clinical characteristic. The change of RPL27 expression occurred in capillaries [49]. Meanwhille, it participate in glucose and lipid metabolism [50]. TXN1P was differentially expressed in Metabolic syndrome, including T2DM [51]. The RPL37 of ribosomal protein genes was the main hub gene in diabetes encephalopathy with well-documented vasoreparative capacity[52–53]. Due to sustained hyperglycemia,microvascular damage was correlated with MNDA [54]. DDX5 were differentially expressed in obese T2DM chronic wounds tissue [55].

When diabetes promotes inflammation, CLEC7A expression may be abnormal [56]. Metformin increase expression of CLEC7A [57]. SIGLEC14 enhanced TNF-alpha secretion and IL-1β release may play a role in inflammation. This effect is related to lipopolysaccharides and NLRP3 inflammasome [58–59]. In WGCNA network interaction results, TNFRSF1A is the most core genes. It is associated with loss of diabetic kidney disease. TNFRSF1A is a positive correlation with HbA1c[60]. Base T2DM is a chronic inflammatory disease, hyperglycemia and NFKB1A were is closely connected[61].

Using the T2D Knowledge Portal, we screened six genes with high expression in T2DM patients. Among them, the GIMAP7, HLA-DQB1 and RPL37 genes are related to triglyceride levels in individuals without T2DM. The study showed that the causal association of triglycerides with diabetes is more obvious in young, middle-aged and nonobese people with T2DM [63]. Although the study investigated the relationship between HLA-DQB1 and type 1 diabetes risk [64], HLA-DQB1 was also shown to be associated with susceptibility and protection of T2DM patients [65]. The genetic characteristics of type 1 diabetes and T2DM might be related to common HLA targets. HLA-DRB5 is related to T2DM, HbA1c and diabetic retinopathy, and the downregulation of HLA-DRB5 expression was associated with an increased risk of T2DM [66]. RPL12 is related to fasting glucose levels and BMI. RPS10 is related to HbA1c and diabetic retinopathy. XIST, HLA-DQA2 and CXCL8 are common DEGs between monocytes and T cells. HLA-DQA2 is related to insulin-like growth factor and neuropathy in T2DM patients, especially in individuals without T2DM [67]. HALLMARK_INTERFERON_GAMMA_RESPONSE and HALLMARK_TNFA_SIGNALING_VIA_NFKB are closely associated with the oxidative stress response [68–69]. These pathways were significantly influenced by the occurrence and development of T2DM. Metformin attenuated inflammatory response and enhanced expressions of neurotrophic factors, thereby protecting OLs via AMPK activation in mixed glial cultures stimulated with lipopolysaccharide/interferon γ in vitro, as evidenced by analysis of the expression of signatory genes of O1(+)/MBP(+) OLs and their cellular populations[70]. Inflammatory cytokines involved in the TNF signalling pathway regulate the insulin signalling pathway through serine phosphorylation to improve T2DM [71]. In addition, T cell receptor signaling pathway may as a pathological mechanism for GDM[72], T2DM phenotype of GK rats may be closely related T cell receptor signaling pathway[73]. The studies reveal that NF-kappa B signaling pathway is a factor involved in the pathobiology of T2DM[74]. Metformin-mediated anti-inflammatory role in macrophages alleviate T2DM[75]. Chemokine signaling pathway has been involved in islet β cell damage[76]. It will be influence the onset and progression of T2DM[77].

Analysis of the pseudotime trajectory of CD4+ T cells and monocytes/macrophages revealed that 50 genes whose expression varied with developmental time were divided into four clusters associated with lymphocyte-mediated immunity, protein folding, immunoglobulins and cytoplasmic translation. In T2DM patients, vascular calcification has been associated with increased S100A9 expression, which promotes the release of extracellular vesicles with a high propensity for calcification from macrophages [78]. Islets under hyperglycaemic conditions trigger an inflammatory response associated with increased expression of S100A8 [79]. Research has shown that plasma S100A12 levels are higher in patients with T2DM than in patients without diabetes. S100A12 may be involved in chronic inflammation in T2DM patients according to stepwise multiple regression analyses [80]. SH3BGRL3 is closely related to insulin-like growth factor (IGF-1). IGF-1 effectively stimulates glucose uptake into muscle tissue and increases glucose metabolism throughout the body, so IGF-1 can lower blood glucose levels by reducing insulin resistance [81]. RPS10 has been shown to be a risk factor for paediatric-onset T2DM, and RPS10 expression was associated with the mother’s allele [82].

The results of intercellular communication analysis showed that CD4+ T cells, CD8+ T cells, NK cells, monocytes and macrophages were in direct and intense communication. CD4+ T cells are helper T lymphocytes that enhance phagocytosis, and CD8+ T cells are inhibitory T lymphocytes that kill target cells. The maintenance of normal immune function depends on the relative stability of the CD4/CD8 ratio. In this study, the number of CD4+ and CD8+ cells in the T2DM group showed an increasing trend, but there was no significant difference from that in the healthy group. The expansion of CD4+ and CD8+ cell populations may contribute to inflammation in the body [83]. Moreover, there was no significant difference in the CD4/CD8 ratio. This finding is consistent with the findings of Miya A et al. They observed no differences in the CD4+ or CD8+ cell numbers between the T2DM group and the non-T2DM group using a blood test on an empty stomach [84].

The interaction between monocytes and macrophages is the closest, and these cells are phagocytes whose main function is to carry out bacteriophage activity and activate immune cells depending on the microenvironment and signalling pathway [85]. T cells and monocytes/macrophages form an extremely sophisticated, complex and well-developed immune system [86].

Macrophage migration inhibitory factor (MIF) is a cytokine that functions as a multipotent inflammatory mediator [87]. When MIF binds to CD74+ and CD44+ cells, it phosphorylates serine, which activates the SRC tyrosine kinase pathway and mediates Akt, AMPK, ERK1, etc., signal transmission [88–89]. MIF-CD74+CD44 mediates intercellular communication and plays important roles in diabetes. MIF can be synthesized by islet β cells and released into the blood as granules that are commonly released with insulin, thus promoting the release of insulin and lowering blood glucose levels [90]. At the molecular level, MIF increases the concentration of GLUT4 and PFK2, which promotes glucose uptake and utilization in skeletal and cardiac muscles, thereby regulating blood glucose levels [91–92]. The expression levels of LGALS9 and CD45 are significantly increased in autoimmune diseases, which is consistent with the results of this study [93].

In conclusion, our transcriptional map of immune cells from PBMCs provides a framework for understanding the immune status of T2DM patients with treatment of metformin via scRNA-seq analysis. In addition, we explored the immune state of T cells and monocytes/macrophages from many perspectives, including functional enrichment, cell differentiation trajectory, and intercellular communication. Analysis of the target genes revealed that they were differentially expressed in each of the two groups, suggesting that new key genes are involved in the metformin treatment T2DM. These factors are all potentially important in the pathogenesis and development of T2DM immunity in PBMCs. Our study also has limitations that should be noted. These results need to be validated by further large scale clinical experiments. Further studies should also include patients who do not respond well to metformin therapy for further comparison. Molecular biology experiments will be performed to validate the mechanisms of the genes. We will also continue our research through numerous experiments.

## Methods

### Experimental samples

A cross-sectional study was used for these analyses. Three healthy samples and five samples from patients with T2DM were included. The Chinese clinical trial registration number is ChiCTR2100049613, and the ethics approval number of the First Affiliated Hospital of Anhui University of Traditional Chinese Medicine is 2021AH-39.

### Diagnostic criteria

Patients were diagnosed with type 2 diabetes mellitus according to the Guidelines for the Prevention and Treatment of Type 2 Diabetes in China. The diagnostic criterion for healthy subjects was a history of systemic disease, such as hypertension, T2DM, cardiopulmonary insufficiency, or the use of other systemic or topical medications, as opposed to patients with T2DM.

The inclusion criterion was age 18-70 years, regardless of sex. These findings are in line with the diagnostic criteria of the healthy population. In line with the diagnostic criteria for T2DM, the patients only received metformin hydrochloride tablets (0.5 g/time, 3 times/day, not less than 3 months; HbA1c ≤ 7.0%). The subjects were informed and voluntarily signed the informed consent form.

The exclusion criteria for patients were type 1 diabetes mellitus, gestational diabetes mellitus, type 2 diabetes mellitus (T2DM) requiring insulin therapy and other special types of diabetes mellitus. Patients with acute complications of diabetes mellitus. Patients with severe cardiovascular and cerebrovascular diseases and severe primary diseases, such as liver, kidney and haematopoietic system diseases. Patients who were allergic to the known ingredients of the study drug and had an allergic constitution. Pregnant and lactating women and those who had recently planned to give birth. Patients with long-term alcoholism, drug dependence or mental illness. Those who had participated in or were participating in other drug clinical trials within one month before the screening period for this study. Those who were not suitable to participate in this clinical study were excluded based on the opinion of the investigator.

### Observation and evaluation indices

Background information included age, sex, height, weight, etc. Safety indicators included vital signs, such as blood pressure, respiration, heart rate, pulse, routine blood, routine urine, ECG, ALT, AST, BUN, CRE, etc. Adverse events, including gastrointestinal symptoms, bleeding, allergy, etc., were recorded, including the time, severity, frequency, duration, measures taken and outcome of adverse events. Efficacy indicators included fasting blood glucose and HbA1c levels.

### Single-cell mRNA sequencing

Afterwards, 2 mL of whole blood containing EDTA was added to a 15 mL centrifuge tube with 3 mL of Ficoll lymphocyte separation media. An equal volume of 1X PBS was added to the blood. Carefully, the diluted blood samples were layered in Ficoll lymphocyte separation liquid. Then, the sample was centrifuged at 400 × g from 18 to 20 ℃ for 30 mins continuously. The mononuclear cell layer was transferred to a 15 mL sterile centrifuge tube using a sterile pipette. Three volumes of 1× PBS were added to the lymphocyte layer, which was carefully mixed by pipetting up and down. Again, the samples were centrifuged at 400 × g, after which the supernatant was discarded. Then, 6 mL of 1×PBS was added to the lymphocyte layer, which was carefully mixed by pipetting up and down. Subsequently, the mixture was centrifuged at 400 × g for 10 mins, after which the supernatant was discarded. The cells were resuspended in the desired volume of 1× PBS. The cellular suspension was stained with 0.4% trypan blue. Cells with greater than 80% viability were qualified for the library construction process. The prepared single-cell suspensions were subsequently partitioned into GEMs (Gel Beads in Emulsions) in an automated Chromium Controller, after which the mRNAs were reverse transcribed into cDNAs. The reaction system was configured in sequence for breaking GEMs, cDNA amplification, fragmentation, end repair, A-tailing, and adaptor ligation PCR. After reacting at a suitable temperature for a fixed period, the products were separately purified. The reaction system was configured so that the products were purified. After library QC, single-stranded PCR products were produced via denaturation. Single-stranded cyclized products were produced by a circularization reaction system. Single-stranded circular DNA molecules were replicated, and a DNA nanoball (DNB) that contained multiple copies of DNA was generated. Sufficient quality DNBs were then loaded into patterned nanoarrays and sequenced through combinatorial probe-anchor synthesis (cPAS).

### Quality control of the single-cell data

The raw gene expression matrix generated from each sample was aggregated using Cell Ranger (v5.0.1) [94], which is provided on the 10x Genomics website. Downstream analysis was performed using the R package Seurat (v 3.2.0) [95]. Specifically, cells with fewer than 200 genes or with > 90% of the proportion of the maximum genes were filtered. For the mitochondrial metric, the cells were sorted in descending order of the mitochondrial read ratio, and the top 15% of cells were filtered. Potential doublets were identified and removed by doublet detection [96]. Cell cycle analysis was performed by using the cell cycle scoring function of the Seurat program. The gene expression dataset was normalized, and subsequent principal component analysis was conducted using only the 2000 highly variable genes in the dataset. UMAP was subsequently used for two-dimensional visualization of the resulting clusters. For each cluster, the marker genes were identified using the Find All Markers function as implemented in the Seurat package (logfc. threshold > 0.25, minPct > 0.1 and Padj ≤ 0.05). Then, the clusters were marked as a known cell type by the SCSA method [97]. Differentially expressed genes across different samples were identified using the Find Markers function in Seurat with the parameter ‘logfc’, threshold > 0.25, minPct > 0.1 and Padj ≤ 0.05’. Volcano plots were created with the R package Enhanced Volcano. The threshold for the log fold change was set at 0.2, and that for p values was set at 0.05. GO (three associated integrated databases: UniProt http://ftp.ebi.ac.uk/pub/databases/GO/goa/UNIPROT/goa_uniprot_all.gaf.gz, NCBI’s gene2GO ftp://ftp.ncbi.nih.gov/gene/DATA/gene2go.gz, GO’s official website: ftp://ftp.pir.georgetown.edu/databases/idmapping/idmapping.tb.gz, downloaded in May 2020) analysis and KEGG (V93.0) pathway analysis were performed using phyper, a function of R. KEGG pathways with FDR ≤0.05 were considered significantly enriched. We used Monocle 2 to extract cells, cluster them and construct quasitime differentiation and development trajectories [98–99]. We performed cell-to-cell interaction analysis with the Cell Phone DB (v2.1.4) [100]. Receptor‒ ligand pairs can be downloaded from https://www.cellphonedb.org/downloads, and significant cell‒cell interactions were selected with a p value < 0.05. We used iTALK (v0.1.0) to perform cell‒cell interaction analysis [101].

### Genetic evidence calculator

The T2DKP contains summary data on genetic correlations, genome annotations, bioinformatics results, expertise in T2D and related traits, blood glucose, etc. The Human Genetic Evidence Calculator integrates several kinds of human genetic results to quantify genetic support for the involvement of a gene in a disease or phenotype of interest [102].

### Statistical analysis

The statistical analyses were conducted in SPSS and the R programming language. A P value less than 0.05 was considered to indicate statistical significance.

## Supporting information

Supporting information is available from the author.

## Acknowledgements

The current study was supported by open bidding for selecting the best candidates of Xin’an medicine and modernization of Traditional Chinese Medicine of IHM (2023CXMMTCM024, 2023CXMMTCM003), University Scientific Research Projects of Anhui (2023AH050782), the National Natural Science Foundation of China (82174153), Anhui University Collaborative Innovation Project (GXXT-2020-025), and Open Fund for Key Laboratory of Glycolipid Metabolism Disease of the Ministry of Education (GYDKFXM01). The authors thank the researchers who provided the original data of open single-cell datasets. In addition, the research groups played an important role and received technical assistance from BGI Genomics.

## Author contributions

Zhao-Hui Fang and Jin-Dong Zhao participated in the design of the study and wrote the manuscript. All the authors have read and approved the final manuscript.

## Data availability statement

The data used to support the findings of this study are included within the article. All the data are available from the corresponding authors or first author upon reasonable request.

## Institutional and use committee statement

All the experiments were performed in accordance with the guidelines approved by the Ethics Committee of the First Affiliated Hospital of Anhui Chinese Medicine University. (Hefei, China, 2023-AH-35)

## REFERENCES

[1] Chiou J, Geusz RJ, Okino ML, et al. Interpreting type 1 diabetes risk with genetics and single-cell epigenomics. Nature. 2021;594(7863):398–402.

[2] Onengut-Gumuscu S, Chen WM, Burren O, et al. Fine mapping of type 1 diabetes susceptibility loci and evidence for colocalization of causal variants with lymphoid gene enhancers. Nat Genet. 2015;47(4):381–386.

[3] Gheibi S, Singh T, da Cunha JPMCM, et al. Insulin/Glucose-Responsive Cells Derived from Induced Pluripotent Stem Cells: Disease Modeling and Treatment of Diabetes. Cells. 2020;9(11):2465.

[4] Chen C, Xiang Q, Liu W, et al. Co-expression Network Revealed Roles of RNA m^6^A Methylation in Human β-Cell of Type 2 Diabetes Mellitus. Front Cell Dev Biol. 2021;9:651142.

[5] Leete P, Oram RA, McDonald TJ, et al. Studies of insulin and proinsulin in pancreas and serum support the existence of aetiopathological endotypes of type 1 diabetes associated with age at diagnosis. Diabetologia. 2020;63(6):1258–1267.

[6] 5Hanna SJ, Powell WE, Long AE, et al. Slow progressors to type 1 diabetes lose islet autoantibodies over time, have few islet antigen-specific CD8^+^ T cells and exhibit a distinct CD95^hi^ B cell phenotype. Diabetologia. 2020;63(6):1174–1185.

[7] Prasad M, Chen EW, Toh SA, Gascoigne NRJ. Autoimmune responses and inflammation in type 2 diabetes. J Leukoc Biol. 2020;107(5):739–748.

[8] SantaCruz-Calvo S, Bharath L, Pugh G, et al. Adaptive immune cells shape obesity-associated type 2 diabetes mellitus and less prominent comorbidities. Nat Rev Endocrinol. 2022;18(1):23–42.

[9] Song Y, He C, Jiang Y, et al. Bulk and single-cell transcriptome analyses of islet tissue unravel gene signatures associated with pyroptosis and immune infiltration in type 2 diabetes. Front Endocrinol (Lausanne). 2023;14:1132194.

[10] Hanna SJ, Tatovic D, Thayer TC, et al. Insights From Single Cell RNA Sequencing Into the Immunology of Type 1 Diabetes-Cell Phenotypes and Antigen Specificity. Front Immunol. 2021;12:751701.

[11] Rai V, Quang DX, Erdos MR, et al. Single-cell ATAC-Seq in human pancreatic islets and deep learning upscaling of rare cells reveals cell-specific type 2 diabetes regulatory signatures. Mol Metab. 2020;32:109–121.

[12] Liu G, Li Y, Zhang T, et al. Single-cell RNA Sequencing Reveals Sexually Dimorphic Transcriptome and Type 2 Diabetes Genes in Mouse Islet β Cells. Genomics Proteomics Bioinformatics. 2021;19(3):408–422.

[13] Huang Y, Cai L, Liu X, et al. Exploring biomarkers and transcriptional factors in type 2 diabetes by comprehensive bioinformatics analysis on RNA-Seq and scRNA-Seq data. Ann Transl Med. 2022;10(18):1017.

[14] Wu H, Gonzalez Villalobos R, Yao X, et al. Mapping the single-cell transcriptomic response of murine diabetic kidney disease to therapies. Cell Metab. 2022;34(7):1064–1078.e6.

[15] Niu T, Fang J, Shi X, et al. Pathogenesis Study Based on High-Throughput Single-Cell Sequencing Analysis Reveals Novel Transcriptional Landscape and Heterogeneity of Retinal Cells in Type 2 Diabetic Mice. Diabetes. 2021;70(5):1185–1197.

[16] Agrafioti P, Morin-Baxter J, Tanagala KKK, et al. Decoding the role of macrophages in periodontitis and type 2 diabetes using single-cell RNA-sequencing. FASEB J. 2022;36(2):e22136.

[17] Elgamal RM, Kudtarkar P, Melton RL, et al. An integrated map of cell type-specific gene expression in pancreatic islets [published online ahead of print, 2023 Aug 15]. Diabetes. 2023;db230130.

[18] Li C, Zhu J, Wei S, et al. Intermittent protein restriction improves glucose homeostasis in Zucker diabetic fatty rats and single-cell sequencing reveals distinct changes in β cells. J Nutr Biochem. 2023;114:109275.

[19] Donath MY, Dinarello CA, Mandrup-Poulsen T. Targeting innate immune mediators in type 1 and type 2 diabetes. Nat Rev Immunol. 2019;19(12):734–746.

[20] Prasad M, Chen EW, Toh SA, Gascoigne NRJ. Autoimmune responses and inflammation in type 2 diabetes. J Leukoc Biol. 2020;107(5):739–748.

[21] Costanzo MC, von Grotthuss M, Massung J, et al. The Type 2 Diabetes Knowledge Portal: An open access genetic resource dedicated to type 2 diabetes and related traits. Cell Metab. 2023;35(4):695–710.

[22] Maiese K. FoxO Transcription Factors and Regenerative Pathways in Diabetes Mellitus. Curr Neurovasc Res. 2015;12(4):404–413.

[23] Arunachalam G, Samuel SM, Marei I, et al. Metformin modulates hyperglycaemia-induced endothelial senescence and apoptosis through SIRT1. Br J Pharmacol. 2014;171(2):523–535.

[24] Kubota N, Kubota T, Itoh S, et al. Dynamic functional relay between insulin receptor substrate 1 and 2 in hepatic insulin signaling during fasting and feeding. Cell Metab. 2008;8(1):49–64.

[25] Kubota T, Kubota N, Kadowaki T. Imbalanced Insulin Actions in Obesity and Type 2 Diabetes: Key Mouse Models of Insulin Signaling Pathway. Cell Metab. 2017;25(4):797–810.

[26] Ustinova M, Ansone L, Silamikelis I, et al. Whole-blood transcriptome profiling reveals signatures of metformin and its therapeutic response. PLoS One. 2020;15(8):e0237400.

[27] Gonzalez FJ, Xie C, Jiang C. The role of hypoxia-inducible factors in metabolic diseases. Nat Rev Endocrinol. 2018;15(1):21–32.

[28] Li L, Pan Z, Yang X. Key genes and co-expression network analysis in the livers of type 2 diabetes patients. J Diabetes Investig. 2019;10(4):951–962.

[29] Guimarães TA, Farias LC, Santos ES, et al. Metformin increases PDH and suppresses HIF-1α under hypoxic conditions and induces cell death in oral squamous cell carcinoma. Oncotarget. 2016;7(34):55057–55068.

[30] Zhou Y, Zhang H, Yao Y, et al. CD4^+^ T cell activation and inflammation in NASH-related fibrosis. Front Immunol. 2022;13:967410.

[31] Berbudi A, Rahmadika N, Tjahjadi AI, et al. Type 2 Diabetes and its Impact on the Immune System. Curr Diabetes Rev. 2020;16(5):442–449.

[32] Touch S, Clément K, André S. T Cell Populations and Functions Are Altered in Human Obesity and Type 2 Diabetes. Curr Diab Rep. 2017;17(9):81.

[33] Liang C, Yang KY, Chan VW, et al. CD8^+^ T-cell plasticity regulates vascular regeneration in type-2 diabetes. Theranostics. 2020;10(9):4217–4232.

[34] Raphael I, Nalawade S, Eagar TN, et al. T cell subsets and their signature cytokines in autoimmune and inflammatory diseases. Cytokine. 2015;74(1):5–17.

[35] Rattik S, Engelbertsen D, Wigren M, et al. Elevated circulating effector memory T cells but similar levels of regulatory T cells in patients with type 2 diabetes mellitus and cardiovascular disease. Diab Vasc Dis Res. 2019;16(3):270–280.

[36] Vivek S, Crimmins EM, Prizment AE, et al. Age-related Differences in T-cell Subsets and Markers of Subclinical Inflammation in Aging Are Independently Associated With Type 2 Diabetes in the Health and Retirement Study [published online ahead of print, 2023 Jun 1]. Can J Diabetes. 2023;S1499-2671(23)00136–3.

[37] Qin W, Sun L, Dong M, et al. Regulatory T Cells and Diabetes Mellitus. Hum Gene Ther. 2021;32(17-18):875–881.

[38] Nishimura S, Manabe I, Nagasaki M, et al. CD8+ effector T cells contribute to macrophage recruitment and adipose tissue inflammation in obesity. Nat Med. 2009;15(8):914–920.

[39] Blériot C, Dalmas É, Ginhoux F, Venteclef N. Inflammatory and immune etiology of type 2 diabetes. Trends Immunol. 2023;44(2):101–109.

[40] Ratter-Rieck JM, Maalmi H, Trenkamp S, et al. Leukocyte Counts and T-Cell Frequencies Differ Between Novel Subgroups of Diabetes and Are Associated With Metabolic Parameters and Biomarkers of Inflammation. Diabetes. 2021;70(11):2652–2662.

[41] Boutens L, Stienstra R. Adipose tissue macrophages: going off track during obesity. Diabetologia. 2016;59(5):879–894.

[42] Blériot C, Barreby E, Dunsmore G, et al. A subset of Kupffer cells regulates metabolism through the expression of CD36. Immunity. 2021;54(9):2101–2116.

[43] Tsay A, Wang JC. The Role of PIK3R1 in Metabolic Function and Insulin Sensitivity. Int J Mol Sci. 2023;24(16):12665.

[44] Del Bosque-Plata L, Martínez-Martínez E, Espinoza-Camacho MÁ, Gragnoli C. The Role of TCF7L2 in Type 2 Diabetes. Diabetes. 2021;70(6):1220–1228.

[45] Guasch-Ferré M, Santos JL, Martínez-González MA, et al. Glycolysis/gluconeogenesis- and tricarboxylic acid cycle-related metabolites, Mediterranean diet, and type 2 diabetes. Am J Clin Nutr. 2020;111(4):835–844.

[46] Matsuura Y, Shimizu-Albergine M, Barnhart S, et al. Diabetes Suppresses Glucose Uptake and Glycolysis in Macrophages. Circ Res. 2022;130(5):779–781.

[47] Ye H, Wang R, Wei J, et al. Bioinformatics Analysis Identifies Potential Ferroptosis Key Gene in Type 2 Diabetic Islet Dysfunction. Front Endocrinol (Lausanne). 2022;13:904312.

[48] Kone M, Pullen TJ, Sun G, et al. LKB1 and AMPK differentially regulate pancreatic β-cell identity. FASEB J. 2014;28(11):4972–4985.

[49] Morgenstern R, Zhang J, Johansson K. Microsomal Glutathione Transferase 1: Mechanism and Functional Roles. Drug Metab Rev, 2011,43(2):300–306.

[50] Suzuki M, Tezuka K, Handa T, et al. Upregulation of ribosome complexes at the blood-brain barrier in Alzheimer’s disease patients. J Cereb Blood Flow Metab. 2022;42(11):2134–2150.

[51] Senevirathna JDM, Yonezawa R, Saka T, et al. Selection of a reference gene for studies on lipid-related aquatic adaptations of toothed whales (Grampus griseus). Ecol Evol. 2021;11(23):17142–17159.

[52] Chitrala KN, Hernandez DG, Nalls MA, et al. Race-specific alterations in DNA methylation among middle-aged African Americans and Whites with metabolic syndrome. Epigenetics. 2020;15(5):462–482.

[53] Wei W, Zhang Q, Jin T, et al. Quantitative Proteomics Characterization of the Effect and Mechanism of Trichostatin A on the Hippocampus of Type II Diabetic Mice. Cell Mol Neurobiol. 2023;43(8):4309–4332.

[54] McLoughlin KJ, Pedrini E, MacMahon M, Guduric-Fuchs J, Medina RJ. Selection of a Real-Time PCR Housekeeping Gene Panel in Human Endothelial Colony Forming Cells for Cellular Senescence Studies. Front Med (Lausanne). 2019;6:33.

[55] Friedrichs P, Schlotterer A, Sticht C, et al. Hyperglycaemic memory affects the neurovascular unit of the retina in a diabetic mouse model. Diabetologia. 2017;60(7):1354–1358.

[56] Boodhoo K, Vlok M, Tabb DL, Myburgh KH, van de Vyver M. Dysregulated healing responses in diabetic wounds occur in the early stages postinjury. J Mol Endocrinol. 2021;66(2):141–155.

[57] Iannucci J, Rao HV, Grammas P. High Glucose and Hypoxia-Mediated Damage to Human Brain Microvessel Endothelial Cells Induces an Altered, Pro-Inflammatory Phenotype in BV-2 Microglia In Vitro. Cell Mol Neurobiol. 2022;42(4):985–996.

[58] Arefin A, Gage MC. Metformin, Empagliflozin, and Their Combination Modulate Ex-Vivo Macrophage Inflammatory Gene Expression. Int J Mol Sci. 2023;24(5):4785.

[59] Yamanaka M, Kato Y, Angata T, Narimatsu H. Deletion polymorphism of SIGLEC14 and its functional implications. Glycobiology. 2009;19(8):841–846.

[60] Tsai CM, Riestra AM, Ali SR, et al. Siglec-14 Enhances NLRP3-Inflammasome Activation in Macrophages. J Innate Immun. 2020;12(4):333–343.

[61] Gómez-Banoy N, Cuevas V, Higuita A, Aranzález LH, Mockus I. Soluble tumor necrosis factor receptor 1 is associated with diminished estimated glomerular filtration rate in colombian patients with type 2 diabetes. J Diabetes Complications. 2016;30(5):852–857.

[62] Seidi A, Mirzaahmadi S, Mahmoodi K, Soleiman-Soltanpour M. The association between NFKB1 -94ATTG ins/del and NFKB1A 826C/T genetic variations and coronary artery disease risk. Mol Biol Res Commun. 2018;7(1):17–24.

[63] Wang X, Liu J, Cheng Z, et al. Triglyceride glucose-body mass index and the risk of diabetes: a general population-based cohort study. Lipids Health Dis. 2021;20(1):99.

[64] Zhang M, Lin S, Yuan X, et al. HLA-DQB1 and HLA-DRB1 Variants Confer Susceptibility to Latent Autoimmune Diabetes in Adults: Relative Predispositional Effects among Allele Groups. Genes (Basel). 2019;10(9):710.

[65] Chinniah R, Vijayan M, Sivanadham R, et al. Association of HLA-A, B, DRB1* and DQB1* alleles and haplotypes in south Indian T2DM patients. Gene. 2016;592(1):200–208.

[66] Jacobi T, Massier L, Klöting N, et al. HLA Class II Allele Analyses Implicate Common Genetic Components in Type 1 and Non-Insulin-Treated Type 2 Diabetes. J Clin Endocrinol Metab. 2020;105(3):dgaa027.

[67] elvaraj MS, Paruchuri K, Haidermota S, et al. Genome-wide discovery for diabetes-dependent triglycerides-associated loci. PLoS One. 2022;17(10):e0275934.

[68] Zheng M, Hu Y, Liu O, et al. Oxidative Stress Response Biomarkers of Ovarian Cancer Based on Single-Cell and Bulk RNA Sequencing. Oxid Med Cell Longev. 2023;2023:1261039.

[69] Grabež M, Škrbić R, Stojiljković MP, et al. A prospective, randomized, double-blind, placebo-controlled trial of polyphenols on the outcomes of inflammatory factors and oxidative stress in patients with type 2 diabetes mellitus. Rev Cardiovasc Med. 2022;23(2):57.

[70] Paintlia AS, Paintlia MK, Mohan S, Singh AK, Singh I. AMP-activated protein kinase signaling protects oligodendrocytes that restore central nervous system functions in an experimental autoimmune encephalomyelitis model. Am J Pathol. 2013;183(2):526–541.

[71] He J, Dai P, Liu L, et al. The effect of short-term intensive insulin therapy on inflammatory cytokines in patients with newly diagnosed type 2 diabetes. J Diabetes. 2022;14(3):192–204.

[72] Chen YM, Zhu Q, Cai J, et al. Upregulation of T Cell Receptor Signaling Pathway Components in Gestational Diabetes Mellitus Patients: Joint Analysis of mRNA and circRNA Expression Profiles. Front Endocrinol (Lausanne). 2022;12:774608.

[73] Liu T, Li H, Ding G, et al. Comparative Genome of GK and Wistar Rats Reveals Genetic Basis of Type 2 Diabetes. PLoS One. 2015;10(11):e0141859.

[74] Bhardwaj R, Singh BP, Sandhu N, et al. Probiotic mediated NF-κB regulation for prospective management of type 2 diabetes. Mol Biol Rep. 2020;47(3):2301–2313.

[75] Xiong W, Sun KY, Zhu Y, Zhang X, Zhou YH, Zou X. Metformin alleviates inflammation through suppressing FASN-dependent palmitoylation of Akt. Cell Death Dis. 2021;12(10):934.

[76] Collier JJ, Sparer TE, Karlstad MD, Burke SJ. Pancreatic islet inflammation: an emerging role for chemokines. J Mol Endocrinol. 2017;59(1):R33–R46.

[77] Qi X, Xing Y, Wang X. Blockade of CCL2/CCR2 Signaling Pathway Exerts Anti-Inflammatory Effects and Attenuates Gestational Diabetes Mellitus in a Genetic Mice Model. Horm Metab Res. 2021;53(1):56–62.

[78] Kawakami R, Katsuki S, Travers R, et al. S100A9-RAGE Axis Accelerates Formation of Macrophage-Mediated Extracellular Vesicle Microcalcification in Diabetes Mellitus. Arterioscler Thromb Vasc Biol. 2020;40(8):1838–1853.

[79] Miyashita D, Inoue R, Tsuno T, et al. Protective effects of S100A8 on sepsis mortality: Links to sepsis risk in obesity and diabetes. iScience. 2022;25(12):105662.

[80] Kosaki A, Hasegawa T, Kimura T, et al. Increased plasma S100A12 (EN-RAGE) levels in patients with type 2 diabetes. J Clin Endocrinol Metab. 2004;89(11):5423–5428.

[81] Moses AC. Insulin resistance and type 2 diabetes mellitus: is there a therapeutic role for IGF-1?. Endocr Dev. 2005;9:121–134.

[82] Miranda-Lora AL, Molina-Díaz M, Cruz M, et al. Genetic polymorphisms associated with pediatric-onset type 2 diabetes: A family-based transmission disequilibrium test and case-control study. Pediatr Diabetes. 2019;20(3):239–245.

[83] Touch S, Clément K, André S. T Cell Populations and Functions Are Altered in Human Obesity and Type 2 Diabetes. Curr Diab Rep. 2017;17(9):81.

[84] Miya A, Nakamura A, Miyoshi H, et al. Impact of Glucose Loading on Variations in CD4^+^ and CD8^+^ T Cells in Japanese Participants with or without Type 2 Diabetes. Front Endocrinol (Lausanne). 2018;9:81.

[85] Kuznetsova T, Prange KHM, Glass CK, de Winther MPJ. Transcriptional and epigenetic regulation of macrophages in atherosclerosis. Nat Rev Cardiol. 2020;17(4):216–228.

[86] Liang C, Yang KY, Chan VW, et al. CD8^+^ T-cell plasticity regulates vascular regeneration in type-2 diabetes. Theranostics. 2020;10(9):4217–4232.

[87] Su H, Na N, Zhang X, Zhao Y. The biological function and significance of CD74 in immune diseases. Inflamm Res. 2017;66(3):209–216.

[88] Lue H, Dewor M, Leng L, et al. Activation of the JNK signalling pathway by macrophage migration inhibitory factor (MIF) and dependence on CXCR4 and CD74. Cell Signal. 2011;23(1):135–144.

[89] Kim BS, Pallua N, Bernhagen J, Bucala R. The macrophage migration inhibitory factor protein superfamily in obesity and wound repair. Exp Mol Med. 2015;47(5):e161.

[90] Benigni F, Atsumi T, Calandra T, et al. The proinflammatory mediator macrophage migration inhibitory factor induces glucose catabolism in muscle. J Clin Invest. 2000;106(10):1291–1300.

[91] Atsumi T, Cho Y R, Leng L, et al. The proinflammatory cytokine macrophage migration inhibitory factor regulates glucose metabolism during systemic inflammation. [J].The Journal of Immunology: Official Journal of the American Association of Immunologists, 2007(8):179.

[92] Verschuren L, Kooistra T, Bernhagen J, et al. MIF deficiency reduces chronic inflammation in white adipose tissue and impairs the development of insulin resistance, glucose intolerance, and associated atherosclerotic disease. Circ Res. 2009;105(1):99–107.

[93] Zhang Y, Lee TY. Revealing the Immune Heterogeneity between Systemic Lupus Erythematosus and Rheumatoid Arthritis Based on Multi-Omics Data Analysis. Int J Mol Sci. 2022;23(9):5166.

[94] Zheng GX, Lau BT, Schnall-Levin M, et al. Haplotyping germline and cancer genomes with high-throughput linked-read sequencing. Nat Biotechnol. 2016;34(3):303–311.

[95] Butler A, Hoffman P, Smibert P, et al. Integrating single-cell transcriptomic data across different conditions, technologies, and species. Nat Biotechnol. 2018;36(5):411–420.

[96] See Gayoso, Adam, & Shor, Jonathan. (2018, July 17). DoubletDetection (Version v2.4). Zenodo. 10.5281/zenodo.2678042

[97] Cao Y, Wang X, Peng G. SCSA: A Cell Type Annotation Tool for Single-Cell RNA-seq Data. Front Genet. 2020;11:490.

[98] Qiu X, Hill A, Packer J, Lin D, Ma YA, Trapnell C. Single-cell mRNA quantification and differential analysis with Census. Nat Methods. 2017;14(3):309–315.

[99] Trapnell C, Cacchiarelli D, Grimsby J, et al. The dynamics and regulators of cell fate decisions are revealed by pseudotemporal ordering of single cells. Nat Biotechnol. 2014;32(4):381–386.

[100] Efremova M, Vento-Tormo M, Teichmann S A, et al. CellPhoneDB: inferring cell–cell communication from combined expression of multisubunit ligand–receptor complexes[J]. Nature protocols, 2020, 15(4): 1484–1506.

[101] Wang Y, Wang R, Zhang S, et al. iTALK: an R Package to characterize and illustrate intercellular communication[J]. bioRxiv, 2019: 507871.

[102] Dornbos P, Singh P, Jang DK, et al. Evaluating human genetic support for hypothesized metabolic disease genes. Cell Metab. 2022;34(5):661–666.

